# FilamentID reveals the composition and function of metabolic enzyme polymers during gametogenesis

**DOI:** 10.1101/2024.03.26.586525

**Authors:** Jannik Hugener, Jingwei Xu, Rahel Wettstein, Lydia Ioannidi, Daniel Velikov, Florian Wollweber, Adrian Henggeler, Joao Matos, Martin Pilhofer

## Abstract

Gamete formation and subsequent offspring development often involve extended phases of suspended cellular development or even dormancy. How cells adapt to recover and resume growth remains poorly understood. Here, we visualized budding yeast cells undergoing meiosis by cryo-electron tomography (cryoET) and discovered elaborate filamentous assemblies decorating the nucleus, cytoplasm, and mitochondria. To determine filament composition, we developed a “Filament IDentification” (FilamentID) workflow that combines multiscale cryoET/cryo-electron microscopy (cryoEM) analyses of gently lysed cells. FilamentID identified the mitochondrial filaments as the conserved aldehyde dehydrogenase Ald4^ALDH2^ and the nucleoplasmic/cytoplasmic filaments being composed of acetyl-CoA synthetase Acs1^ACSS2^. The near-native high-resolution structures revealed the mechanism underlying polymerization and enabled us to perturb filament formation. Acs1 polymerization facilitates the recovery of chronologically aged spores, and more generally, the cell cycle re-entry of starved cells. FilamentID is broadly applicable to characterize filaments of unknown identity in diverse cellular contexts.

**HIGHLIGHTS:**

- FilamentID: a multiscale imaging workflow to characterize cellular filaments of unknown composition
- The conserved aldehyde dehydrogenase Ald4^ALDH2^ polymerizes into filament arrays within meiotic mitochondria
- The conserved acetyl-CoA synthetase Acs1^ACSS2^ forms filament arrays in the nucleus and the cytoplasm
- Metabolites mediate Acs1 polymerization to store Acs1 in an inactive state in gametes and starved cells
- Acs1 filament formation is required for efficient return to growth from starvation stress

## INTRODUCTION

Most organisms live in rapidly changing environments and frequently experience adverse conditions for growth and reproduction. In microorganisms, one common response to environmental stress is the reversible entry into a dormant state, in which cells reduce their metabolic activity and arrest proliferation. Dormancy can also involve the execution of specialized developmental programs that result in differentiated cell types, such as spores, which can withstand nutrient scarcity, non-optimal temperatures, or desiccation. The formation of dormant cells typically relies on extensive remodeling of the cellular architecture, including, amongst others, changes in cell size, cell envelope structure, or cytoplasmic density and composition ^1,2^. These ultrastructural adaptations are thought to enable cells to enter, maintain, and recover from dormancy, but their underlying molecular mechanisms remain poorly understood.

A hallmark feature of cellular dormancy is a decrease in metabolic activity ^1^. In addition to the downregulation of enzyme function through the modulation of gene expression, an emerging concept posits that metabolic activity can be spatiotemporally controlled by enzyme self-assembly into filamentous polymers or agglomerates. Subcellular compartmentalization of metabolic enzymes mediated by self-association has been observed in different domains of life and has often been associated with cellular starvation ^3–9^. In the budding yeast *Saccharomyces cerevisiae*, genome-wide fluorescence light microscopy screens have revealed a surprisingly high number of enzymes that form punctate or rod-shaped structures in response to the metabolic state of the cell ^10–13^. Notable examples include the cytidine triphosphate synthase CTP ^14,15^, the glutamine synthetase Gln1 ^16^, the translation initiation factor elF2B ^17^, or the glucokinase Glk1 ^18^. In all four cases, filament formation has been proposed to play important roles in the regulation of enzyme activity and, as such, to be of relevance for cell physiology. Even though biochemical reconstitution approaches coupled with structural studies have begun to shed light on the mechanistic basis of filament organization, it remains challenging to ascertain filament structural organization in the cellular context. Moreover, whether filament formation occurs and plays specialized roles in other biological contexts, for example when cells enter prolonged periods of developmental arrest or dormancy after gametogenesis, remains to be determined.

In many sexually reproducing organisms, including budding yeast, cells enter gametogenesis as a natural response to nutrient deprivation, resulting in the production of highly differentiated spores ^19^. The principal features of meiosis are evolutionarily conserved and include a single round of DNA replication, followed by two consecutive nuclear divisions, termed meiosis I and meiosis II ^20^. At the completion of meiosis, a haploid DNA complement, cytoplasmic content, and organelles are inherited by four arising gametes, which are protected by a thick spore wall. The resistant spores remain viable in a dormant state, preserving the potential to resume growth after encountering favorable environments ^19^. How spores preserve all essential cellular components required for the resumption of growth over potentially very long periods of time remains unclear.

Here, we have used cryo-electron tomography (cryoET) imaging to study budding yeast cells throughout gametogenesis, since it allows the visualization of mesoscale assemblies in their cellular context in a near-native state ^21,22^. We discovered distinct types of previously uncharacterized filament assemblies in different cellular compartments. Since the identification of such structures by cryoET remains exceptionally challenging, we developed a workflow to image gently lysed cells by employing a combination of cryoET and single-particle cryo-electron microscopy (cryoEM), which we have termed FilamentID. By enabling us to obtain high-resolution structures in near-native conditions, FilamentID allowed the unambiguous identification of two distinct types of polymerized metabolic enzymes. Targeted mutations and downstream functional experiments suggest that metabolic enzyme polymerization is induced during starvation and required for efficient gamete awakening after prolonged dormancy.

## RESULTS

### Filament assemblies form in mitochondria, nucleus, and cytoplasm of yeast cells undergoing gametogenesis

We set out to visualize ultrastructural adaptations of cells undergoing gametogenesis upon nutrient starvation. To this end, we synchronized diploid *S. cerevisiae* cells at G0/G1 through gradual starvation and induced meiosis by transferring the cultures into sporulation medium (SPM) lacking a nitrogen source, as previously described (Figure S1A) ^23,24^. Fluorescence activated cell sorting (FACS analysis) of DNA content showed that most cells were in G0/G1 prior to transfer to SPM and efficiently underwent DNA replication between two and four hours after induction of meiosis (Figure S1B). For imaging in a near-native state, meiotic cells were plunge-frozen on EM grids, thinned by cryo-focused ion beam (cryoFIB) milling, and imaged by cryoET ^25^. To reduce cell-to-cell variability that could arise from asynchronous meiotic progression, in an initial step we used prophase I-arrested cultures (*ndt80Δ*) ^26^ that were collected eight hours after induction of meiosis.

We observed various filamentous assemblies in different sub-cellular compartments (Figure 1A-C). One type of filament was frequently observed in mitochondria, hereafter referred to as “mitochondrial filament.” In longitudinal views, these filaments appeared straight and were often bundled together. Cross-sections in the representative cryo-tomogram revealed that the filament bundles were ordered in arrays of ∼40 individual filaments. A second type of filament was found in the nucleus and in the cytoplasm, hereafter referred to as “nucleocytoplasmic” filament. These nucleocytoplasmic filaments were also arranged in filament bundles, however, the organization appeared more heterogeneous and complex, with some filaments branching away from the bundles. Cross-sections of individual nucleocytoplasmic filaments showed a triangular architecture.

**Figure 1.**
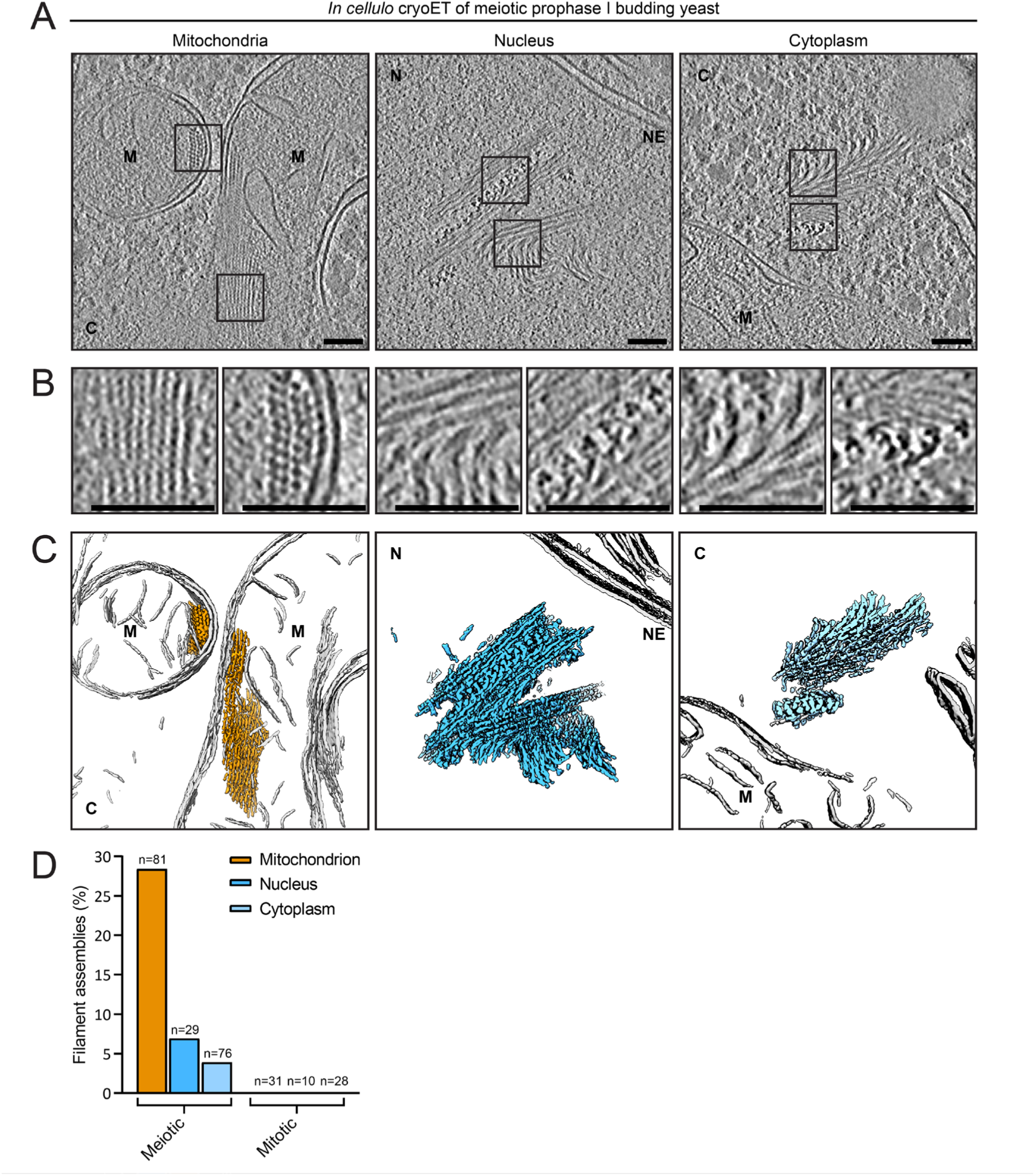
Filament assemblies form in the mitochondria, nucleus, and cytoplasm of meiotic yeast cells. See also Figure S1. **(A)** Slices through cryo-tomograms of FIB-milled meiotic prophase I budding yeast cells with filament assemblies in the mitochondria, in the nucleus, and in the cytoplasm. Shown are projections of 9.14 nm (mitochondria) and 8.68 nm (nucleus and cytoplasm) thick slices. Cells were collected from a *ndt80Δ* strain arrested in prophase I, 8 h post induction of meiosis. M: mitochondrion, C: cytoplasm N: nucleus, NE: nuclear envelope. Scale bars: 100 nm. **(B)** Magnified views of cryo-tomograms shown in (**A**). Note that the top views of single filaments display a round shape for mitochondrial filaments, whereas the nuclear and cytoplasmic filaments have a triangular shape. Scale bars: 100 nm. **(C)** Segmentation models of cryo-tomograms in (**A**) showing filament arrays and bundles. Mitochondrial filaments (orange), nuclear filaments (blue), and cytoplasmic filaments (light blue) are highlighted. Organelle membranes are colored in gray. **(D)** Filament assemblies in the different cellular compartments are observed in meiotic but not in mitotically dividing cells. Meiotic mitochondria (n = 81), nuclei (n = 29), and cytoplasm (n = 76) and mitotic mitochondria (n = 31), nuclei (n = 10), and cytoplasm (n = 28) were analyzed. The fraction (%) of mitochondria, nuclei, and cytoplasm containing the main types of filaments described in (**A-C**) is indicated for each cellular context. Numbers for analyzed compartments (n) represent cumulated results from at least two independent experiments.

Quantification of the two filament types revealed that mitochondrial filaments were present in ∼28% of the imaged mitochondria (n = 23/81), and nucleocytoplasmic filaments in ∼7% of the tomograms containing nuclear volume (n = 2/29) and ∼4% of the tomograms containing cytoplasmic volume (n = 3/76) (Figure 1D). Importantly, no filaments were observed in mitotically dividing cells grown in rich medium, indicating that meiosis, or the process of starvation-induced gametogenesis, may drive the formation of such structures.

### FilamentID: a workflow to identify cellular filaments of unknown molecular composition

Conventional approaches to identify filaments seen *in cellulo* would be either A) the generation of high-resolution sub-tomogram averages, or B) the purification of the filaments followed by downstream single-particle cryoEM analyses. For the identification of filaments, both approaches are very challenging, since the abundance of the filaments is relatively low and their identities are unknown. We therefore developed a “Filament IDentification” (FilamentID) workflow that is applicable to low-abundance targets (problematic for approach “A”) and that circumvents the need for purification of the complex of interest (required for approach “B”). One critical aspect of the workflow entails the preparation of the sample such that it can be imaged without the need for cryo-sample thinning. Towards this goal, we adapted a chromosome surface spreading protocol ^27,28^ to gently lyse and “spread” either isolated organelles or entire cells onto EM grids. Since cell envelope and organelle integrity become partially disrupted, the cellular contents are diluted, making intracellular complexes directly accessible for a combination of cryoET and single-particle cryoEM imaging.

The workflow features two variations for the identification of filaments in mitochondria or nucleoplasm/cytoplasm, respectively (Figure 2). Step 1: Cells are spheroplasted by digesting the cell wall with Zymolyase. For the identification of mitochondrial filaments, mitochondria are then purified and spread onto EM grids by hypo-osmotic swelling. For the identification of nucleocytoplasmic filaments, entire spheroplasts are spread onto an EM grid by hypo-osmotic swelling and the addition of a mild detergent. Step 2: Grids are vitrified by plunge-freezing. Step 3: Samples are subjected to data collection. Each sample is imaged by recording tilt-series for cryoET, as well as 2D projection images for targeted single-particle cryoEM. Step 4: CryoET data is used to generate low-resolution sub-tomogram averages to assess filament symmetry. The single-particle data are used to generate high-resolution maps by helical reconstruction, based on the symmetry information determined by cryoET. Step 5: Integrating the cryoEM data with structure prediction, structure databases, and proteomic analyses allows for the unambiguous identification of the filament components. Step 6: Structural findings are validated by genetically interfering with filament polymerization. The resulting point mutants can be used in functional assays to reveal the role of filament formation. We note that the imaging of entire spread spheroplasts would also be fully compatible with the identification of mitochondrial filaments. However, the mitochondrial purification step is useful in order to reduce the workload in steps 3-5.

**Figure 2.**
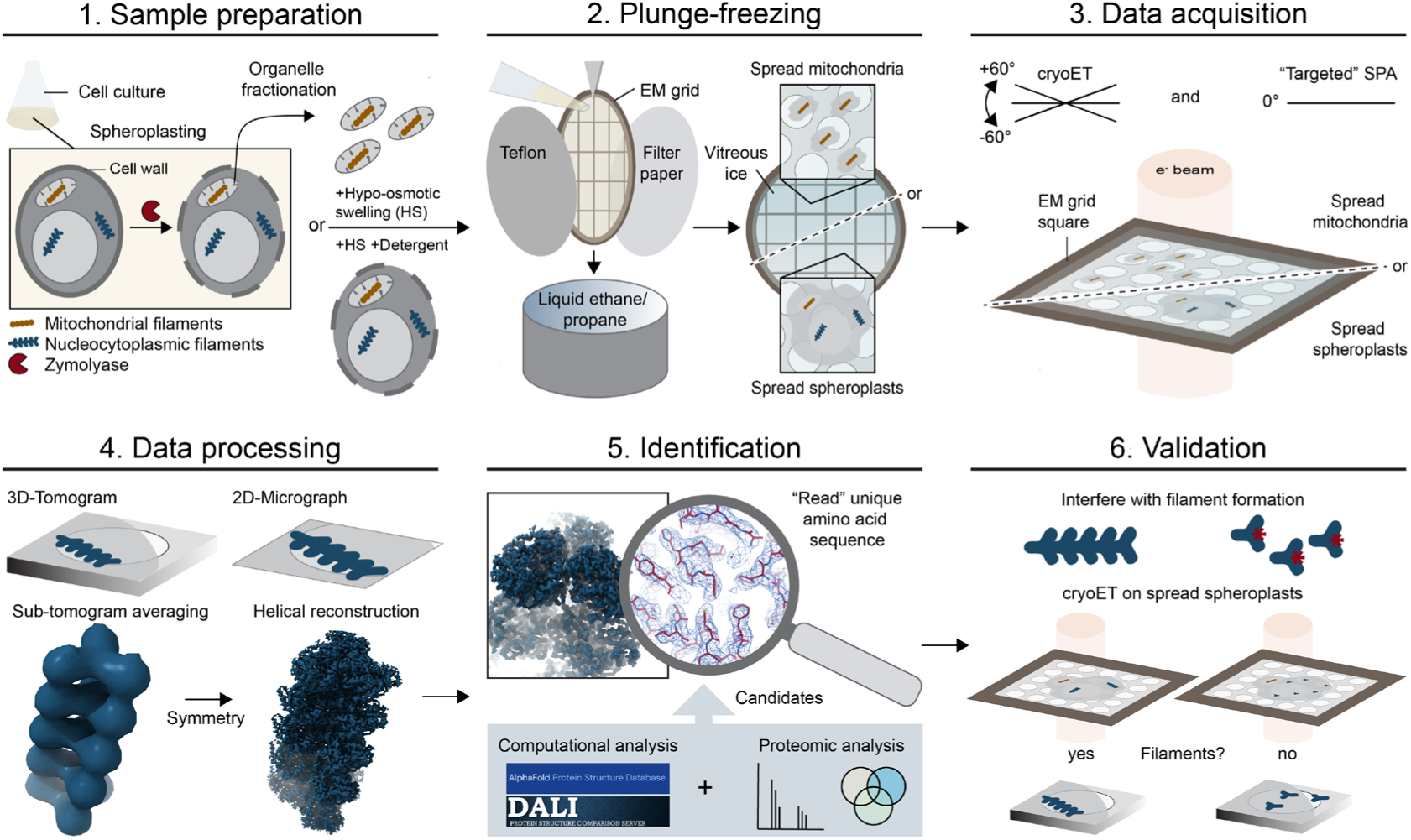
FilamentID: a workflow to identify cellular filaments of unknown molecular composition. See also Figure S2/S5. Schematic workflow to IDentify the composition of Filament assemblies (FilamentID) observed *in cellulo*. Key steps (1-6): (1) Sample preparation: cells are spheroplasted by digesting the cell wall with Zymolyase. Depending on the subcellular localization of the filaments, either mitochondria are purified and spread onto EM grids, or entire cell spheroplasts are spread onto EM grids by hypo-osmotic swelling and the addition of detergent. (2) Plunge-freezing: samples are vitrified by plunge-freezing. (3) Each sample is imaged by recording tilt series for cryoET, as well as 2D projection images for single-particle cryoEM. (4) Data processing: the cryoET data are used to generate sub-tomogram averages to determine filament symmetry. Subsequently, single-particle data are used to generate high-resolution maps by helical reconstruction, based on the initial symmetry information determined by cryoET. (5) Identification: information from computational analysis is used to identify the filament component(s). Depending on the level of resolution obtained, complementary information from proteomics can be used to narrow down candidates. (6) Validation: candidate gene(s) are deleted and cryoET is used to verify the impact on filament structure. In addition, the structural information obtained is used to guide the perturbation of filament formation by the introduction of specific point mutations. Note: it also would have been possible to identify the mitochondrial filaments without the mitochondrial purification in step 1; however, the mitochondrial purification step significantly simplifies steps 3-6. For simplicity, in steps 4-6 only the nucleocytoplasmic filament is shown.

### The conserved mitochondrial aldehyde dehydrogenase Ald4 polymerizes into filaments

We first applied FilamentID to characterize the mitochondrial filaments. To determine whether we could preserve the filament arrays after organelle purification, we imaged isolated mitochondria by cryoET. We observed the characteristic mitochondrial double membrane, cristae decorated with F_O_F_1_-ATP synthase, as well as individual filaments and ordered filament arrays (Figure 3A and S2). Sub-tomogram averaging of the filaments revealed a repetitive two-fold symmetric architecture that matches the *in cellulo* observations (Figure 1A-C and 3B).

**Figure 3.**
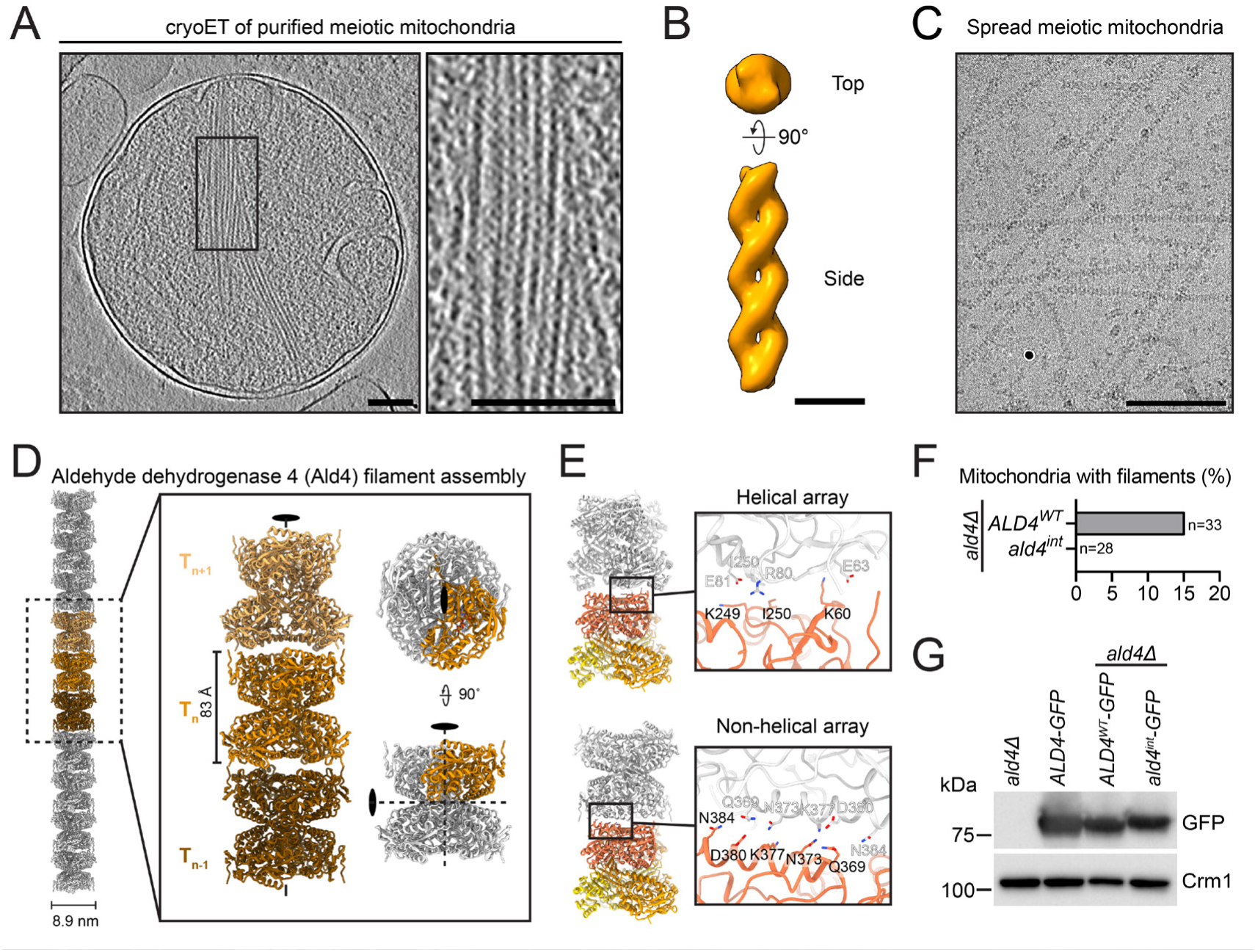
The conserved mitochondrial aldehyde dehydrogenase Ald4 polymerizes into filaments. See also Figure S2-S4. **(A)** Slice through a cryo-tomogram of purified mitochondria. Note that straight filament arrays can cross the entire width of the mitochondrion as depicted. Magnified view of the boxed area is shown on the right. Cells were collected from a *ndt80Δ* strain arrested in prophase I, 6 h post induction of meiosis. Shown are projections of 5.5 nm thick slices. Scale bars: 100 nm. **(B)** Sub-tomogram averaging of filaments from purified meiotic mitochondria shown in (**A**) reveals a two-fold symmetry. Shown are top and side views. Scale bar: 10 nm. **(C)** Purified mitochondria in (**A**) were spread and imaged by cryoEM. A representative micrograph with individual filaments observed in spread meiotic mitochondria is shown. Scale bar: 100 nm. **(D)** Structural model of Ald4 in a filamentous form. Left: stacked array of one Ald4 filament, in which three consecutive Ald4 tetramers are colored in different shades of orange. Right: magnified view of the central three Ald4 tetramers on the left. The top and side views of the ribbon diagram of the central Ald4 tetramer are shown on the right, where a single Ald4 subunit is colored orange. The two-fold axis of Ald4 is represented by an ellipse. **(E)** Ribbon diagrams showing the putative interfaces between two consecutive Ald4 tetramers in helical (top) and non-helical (bottom) arrays. The subunits of one Ald4 tetramer are colored in different shades of orange, while the other Ald4 tetramer is colored white. The detailed interface contacts are shown on the right, where the side chains of putative residues mediating contacts are labeled and shown in stick diagrams. **(F)** The Ald4 interface shown in (**E**) is needed for filament formation. Purified mitochondria from meiotic cultures (6 h after induction of meiosis) of the indicated genotypes were analyzed by cryoET as described in more detail in Figure S4E. Shown are percentages of mitochondria containing filaments and the total number of mitochondria (n) analyzed for each genotype are indicated. **(G)** Ald4 interface mutations do not affect protein stability. Western blot analysis from haploid cell cultures collected after 24 h in YPA media. Ald4-GFP expression levels were probed with anti-GFP antibodies and compared in strains with the indicated genotypes. Crm1 was used as protein normalization control.

Next, we collected single-particle cryoEM data of spread mitochondria. Due to the heterogeneous distribution of the filaments on the grid, we employed a targeted single-particle cryoEM data collection strategy, where we extensively screened grid squares at low magnification, followed by high-magnification data collection (Figure 3C). As detailed below, due to the repetitive architecture of the filaments, a small dataset of several hundred micrographs was sufficient for filament identification. For helical reconstruction, we used the symmetry information obtained from sub-tomogram averaging as a starting point to interpret the helical parameters of the filament in real space (Figure 2). We determined an initial filamentous structure at 6.8 Å resolution (Figure S3A). As the resolution was not sufficient to trace the Cα-backbone, we employed structural dockings of AlphaFold-predicted mitochondrial proteins ^29,30^. To identify potential candidates, we have used mass spectrometry (MS) analysis in a parallel study to characterize the meiotic proteome ^31^, which revealed that various mitochondrial matrix/inner membrane components ^32^ are upregulated at the onset gametogenesis (Figure S4A). Out of 30 candidate proteins, we identified the metabolic enzyme aldehyde dehydrogenase 4 (Ald4) as the most likely filament component (Figure S4B). Based on the knowledge that aldehyde dehydrogenases form tetramers ^33^, we further performed 2D classification and found that the repeating subunits polymerize in two distinct modes, helical and non-helical (Figure S3B). Therefore, we then treated the central part of the helical segments as individual particles and determined the Ald4 tetramer at a final resolution of 3.8 Å (Figure S3C-D). Ald4 tetramers can stack together to form a helical filament architecture (rise = 83 Å, twist = ∼180°), which fitted our map from sub-tomogram averaging (Figure S4C). Moreover, Ald4 tetramers could also assemble in a head-to-head and tail-to-tail fashion (non-helical mode) (Figure 3D and S3B-C).

To validate the presence of Ald4 in the filaments, we first analyzed the reconstructed map and found that it was incompatible with other Ald isoforms present in yeast (Figure S3E, S4D & Table S1). In addition, we generated an *ALD4* deletion mutant (*ald4Δ*), isolated mitochondria from meiotic cells, and imaged them using cryoET. In *ald4Δ* strains, we did not observe any filaments in mitochondria (n = 0/77), whereas ∼14% of the mitochondria from wild-type cells contained filaments (n = 10/71) (Figure S4E).

To understand how Ald4 assembles into filaments, we exploited the tetramer stacking interfaces and identified eight residues that potentially mediate filament polymerization (Figure 3E). We mutated these eight residues and tested if strains expressing the interface mutations (*ald4^int^*) could form filaments in isolated mitochondria. Notably, we did not detect any filaments in the *ald4^int^* mitochondria (n = 0/28), whereas ∼15% of the mitochondria from the *ALD4^WT^* control strain contained filaments (n = 5/33) (Figure 3F). Western blotting showed that the expression levels of GFP-tagged *ALD4^WT^* and the *ald4^int^*mutant were comparable (Figure 3G), suggesting that the interface mutations prevent filament assembly without affecting the general stability of the protein.

In summary, our workflow unambiguously identified the mitochondrial filament arrays as homopolymers of Ald4, and we report the first high-resolution structure of a eukaryotic polymerized aldehyde dehydrogenase enzyme. Our filament identification is consistent with a previous report of elongated inclusion bodies observed by conventional EM, which co-localized with Ald4 antibodies in immuno-gold staining ^34^. An accompanying paper ^31^ presents a characterization of the role of Ald4 polymerization during yeast gametogenesis.

### The conserved acetyl-CoA synthetase Acs1 forms filament assemblies in the nucleus and the cytoplasm

We then applied FilamentID to characterize the nucleocytoplasmic filament assemblies (Figure 1 A-C). To this end, we spread entire meiotic yeast cells directly on EM grids and plunge-froze them (Figure 2). Spread spheroplasts appeared as disc-shaped densities, with ∼4 µm diameter in low magnification images (Figure S5A). Cryo-tomograms of spreads revealed not only a variety of different filament types, but also other well-preserved molecular superstructures, such as spindle pole bodies connected via microtubules (Figure S5A-B). Interestingly, the nucleocytoplasmic filaments were seen both as individual filaments and organized as complex bundles (Figure 4A). In some of the tomograms we also observed potential (dis-) assembly intermediates, where small filaments appear to converge into a large filament bundle (Figure S5C). Sub-tomogram averaging of individual filaments revealed a triple-helical complex with 5.5 nm ladder-like repeats (Figure 4B). The triangular shape of the averaged volume, as well as the overall bundle organization resembled our initial observations *in cellulo* (Figure 1A-C).

**Figure 4.**
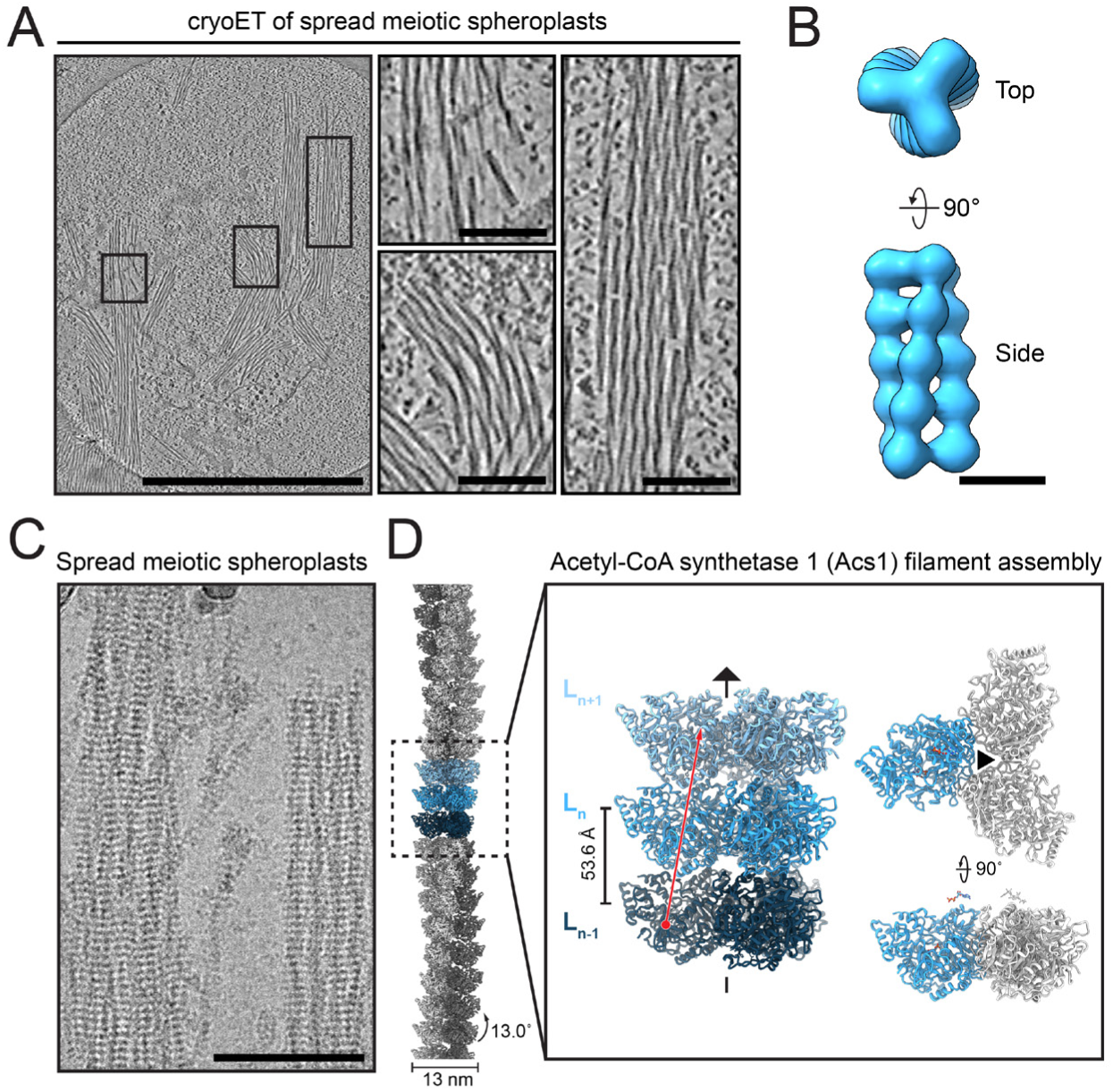
The conserved acetyl-CoA synthetase Acs1 forms filament assemblies in the nucleus and the cytoplasm. See also Figure S5-S7. **(A)** Slice through a cryo-tomogram of spread meiotic yeast spheroplasts enable a detailed view of the nucleocytoplasmic filaments assemblies observed in Figure 1A. Magnified views of the boxed areas highlight single filaments as well as straight and bent bundles. Cells were collected from a *ndt80Δ* strain arrested in prophase I, 8 h post induction of meiosis. Shown are projections of 9.14 nm thick slices. Scale bars: 1 µm and 100 nm for magnified views. **(B)** Sub-tomogram averaging of filaments from yeast spheroplasts shown in (**A**) reveals a three-fold symmetry with a ladder-like repeat. Shown are top and side views. Scale bar: 10 nm. **(C)** Representative cryoEM micrograph of filament bundles from spread meiotic yeast spheroplasts as shown in (**A**). Scale bar: 100 nm. **(D)** Structural model of Acs1 in its filamentous form. Left: Helical array of triple-helical Acs1 filament. A single helical strand is highlighted in dark gray and three consecutive layers are colored in different shades of blue. Right: Magnified view of the central three layers on the left. The single helical strand is marked with a red arrowhead. The side and top view of the ribbon diagram of the central Acs1 trimer are shown on the right, where a single Acs1 subunit is colored in blue. The three-fold axis of the Acs1 trimer is represented by a triangle.

To complement the cryoET data, we applied targeted single-particle cryoEM data collection (Figure 4C), followed by helical reconstruction (Figure S6A-B). This allowed us to obtain a filament structure at ∼3.5 Å resolution (Figure S6C-D), which was sufficient to directly trace the Cα-backbone of one subunit. The backbone model was manually built and was then subjected to the Dali server ^35^, which revealed the metabolic enzyme acetyl-coenzyme A synthetase 1 (Acs1) as a strong candidate.

To validate that the filament consists of Acs1 and not Acs2, a second isoform present in yeast, we compared their sequences and analyzed densities of characteristic residues in our atomic model (Figure S6E, Figure S7A & Table S1). In addition, we generated an *ACS1* deletion mutant (*acs1Δ*) and performed cryoET on the spread spheroplasts of meiotic cells. We did not observe any nucleocytoplasmic filament bundles in spreads from *acs1Δ* spheroplasts (n = 0/80), while ∼32% of the wild-type spheroplasts contained filament bundles (n = 20/63) (Figure S7B).

The filament architecture consists of layers of Acs1 trimers, which are related to consecutive layers through a helical operation (rise = 53.61 Å, twist = 13.03°), resulting in a filament with 13 nm diameter (Figure 4D). Structural docking showed that Acs1 filaments had the same arrangements as seen in the sub-tomogram average (Figure S7C), whereas the filament arrangement is different from the packing of yeast Acs1 in the previously resolved crystal structure (PDB entry: 1RY2) (Figure S7D) ^36^.

In summary, we report here the first high-resolution structure of Acs1 homopolymers in a natively polymerized state. These filaments form elaborate bundles in the nucleoplasm and cytoplasm of meiotic yeast cells. Notably, nucleocytoplasmic filament bundles have previously been observed by others in meiotic yeast cells ^37^ and were proposed to be actin-based/associated ^38^. Using our novel FilamentID approach, we characterized filament structure at higher resolution, which enabled the unambiguous identification of the filament building blocks.

### Metabolites mediate Acs1 filament assembly to inhibit acetyl-CoA production

Next, we set out to elucidate the functional relevance of Acs1 polymer formation. The first step towards this goal was to determine the enzymatic state of filamentous Acs1. The metabolic reaction catalyzed by Acs1 consists of two steps (Figure 5A). In the first step, Acs1 produces acetyl-AMP and diphosphate from acetate and ATP. In the second step, the acetyl-AMP intermediate reacts with CoA to generate acetyl-CoA ^39^. During the enzymatic reaction, the large N-terminal domain (NTD) mediates the binding of ATP substrate, whereas the small C-terminal domain (CTD) undergoes a large conformational change to catalyze the second step of the reaction ^36,40^ (Figure 5B).

**Figure 5.**
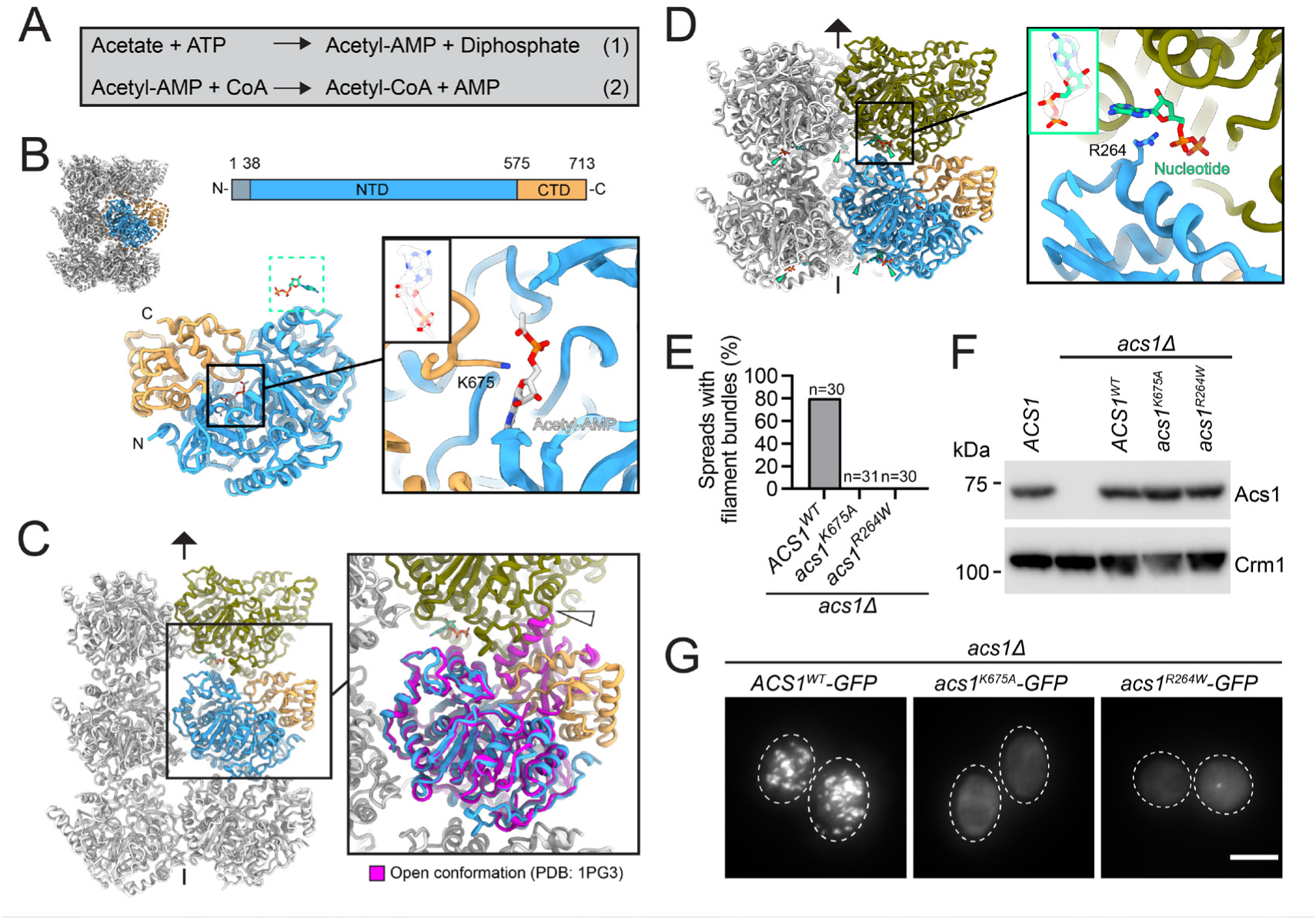
Metabolites mediate Acs1 filament assembly to inhibit acetyl-CoA production. See also Figure S8. **(A)** Acs1 uses acetate and ATP to produce acetyl-CoA in a two-step reaction. In the first step, an acetyl-AMP intermediate is formed, and diphosphate is released. In a second step, acetyl-AMP reacts with CoA to produce acetyl-CoA with the release of AMP ^39^. **(B)** Ribbon and stick diagrams showing two types of metabolites bound to Acs1 in the filament. Top left: the helical array of three Acs1 layers, where one subunit is colored and highlighted with a dashed line. Top right: Scheme showing the overall Acs1 domain organization, where the N-terminal large domain (NTD) and the C-terminal small domain (CTD) are colored blue and orange respectively. The flexible N-terminal part, which is not resolved in the structure, is colored grey. Bottom-left: the structural model of one Acs1 subunit with two metabolites bound (acetyl-AMP: gray; nucleotide (potentially ADP): green). The position of the nucleotide is highlighted with a green dashed box and the details are shown in panel (**D**). Bottom-right: Magnified view showing the contacts between acetyl-AMP and Acs1. The side chain of the residue in the catalytic center (K675) is shown in stick diagram, while the corresponding densities around acetyl-AMP are transparent. **(C)** Ribbon diagrams illustrating the predicted structural clashes that prevent the second step in acetyl-CoA synthesis when Acs1 is in a filamentous form. Left: helical assembly of three Acs1 trimer layers, where one Acs1 subunit is color-coded as in (**B**) and one subunit from the adjacent trimer layer is colored olive. Right: structural superposition of one Acs1 subunit with CoA-bound bacterial ACS (PDB entry: 1PG3) ^40^. Note the severe steric clashes (highlighted with white arrowhead) between the bacterial homolog (magenta) and the Acs1 molecules in the adjacent filament layer (olive). **(D)** Ribbon and stick diagrams showing the metabolic nucleotide (potentially ADP) bridging two Acs1 adjacent layers. Left: the structure of Acs1 subunits from two adjacent layers. The color code of two subunits matches panel (**C**). Right: Magnified view showing the contacts between the nucleotide (green) and Acs1 subunits. The side chain of the residue binding to the nucleotide (R264) is shown in stick diagrams. The corresponding densities around the nucleotide are shown in transparent. **(E)** No filament bundles are detected in point mutation strains of the residues shown in (**B**) and (**D**). Spread meiotic yeast spheroplasts (6 h after induction of meiosis) for the indicated genotypes were analyzed by cryoEM as described in more detail in Figure S7D. Shown are percentages of spreads containing Acs1 filament bundles in EM images and the total number of spreads (n) analyzed for each genotype are indicated. **(F)** Acs1 point mutations do not affect protein stability. Western blot analysis from meiotic cell cultures collected 6 h after induction of meiosis. Acs1 expression levels were probed with an anti-Acs1 antibody and compared in strains with the indicated genotypes. Crm1 was used as protein normalization control. **(G)** Fluorescent rods and foci formation is disrupted in *ACS1* point mutation strains C-terminally tagged with GFP. Shown are representative maximum intensity projections of the Acs1-GFP signal expressed from the endogenous promotor for the indicated genotypes. Cells were collected 6 h after induction of meiosis. Cell walls are indicated with a dashed white line. Scale bar: 5 µm.

We found that Acs1 subunits in filaments have a conformation that is similar to the crystal structure of the binary complex with AMP (PDB entry: 1RY2) (R.M.S.D. 1.30 Å, Figure S8A), which represents the structure of the first step of the enzymatic reaction ^36^. The acetyl-AMP intermediate could be observed in the active site of the NTD and is located close to the catalytic residue K675 (Figure 5B and S8B) ^41^. Moreover, no acetyl-CoA product of the second step of the reaction was found in our reconstructed map. In order to carry out the second step of the reaction, the Acs1 CTD has to undergo a conformational change, which would result in a severe clash with the adjacent trimer layer shown by a structural superposition with the bacterial ACS representing the open conformation of the second step of the reaction (PDB entry: 1PG3; Figure 5C) ^40^. Therefore, we propose that Acs1 in filaments catalyzed the formation of acetyl-AMP. However, once in its filamentous form, Acs1 is sterically hindered by an adjacent Acs1 trimer layer to produce the final product, acetyl-CoA.

To determine if the formation of acetyl-AMP is required for filament assembly, we mutated the key residue K675 in the catalytic site and monitored the presence of filament bundles in spread spheroplasts with cryoET. We did not observe any filament bundles in spreads from *acs1^K^*^675^*^A^* spheroplasts (n = 0/31), while ∼80% of the control strain (*ACS1^WT^*) contained filament bundles (n = 24/30) (Figure 5E). Western blotting using antibodies raised against Acs1 showed comparable protein expression (Figure 5F). Moreover, we complemented this analysis by monitoring the cellular distribution of GFP-tagged Acs1. We observed fluorescent rods and foci in the *ACS1^WT^-GFP* strain but did not detect such structures in the *acs1^K^*^675^*^A^-GFP* mutant (Figure 5G). These data suggest that the catalytic activity of Acs1, in particular the formation of the acetyl-AMP intermediate, is required for filament bundle formation.

Interestingly, we detected three additional densities at the interface between Acs1 trimer layers. These extra densities likely correspond to metabolites, possibly ADP nucleotides (Figure 5D). Each putative nucleotide binds to the surface of α helices in the Acs1 NTDs (Figure S8C). The putative nucleotides are likely further stabilized via the α helix from the consecutive Acs1 NTD, which directly contacts the positively charged residue R264 (Figure 5D). Additional interactions between two consecutive Acs1 trimer layers, especially salt bridges, are likely to contribute further to filament stability (Figure S8D). To explore the role of the putative nucleotide in Acs1 filament assembly, we blocked the corresponding binding site by mutating R264 to the bulky residue tryptophan. Strikingly, we no longer detected elaborate filament bundles in spreads of meiotic *acs1^R^*^264^*^W^* spheroplasts (n = 0/30), even though the expression level of Acs1^R264W^ was comparable to Acs1^WT^ (Figure 5E-F). Additionally, most meiotic cells of *acs1^R^*^264^*^W^-GFP* mutants contained only diffuse fluorescent signal (Figure 5G).

Overall, our structural and mutant analyses indicate that Acs1 polymerization is mediated by two metabolites, acetyl-AMP bound in the catalytic pocket and a putative nucleotide bound at the filament layer interface.

### The catalytic activity of Acs1 is required for sporulation and germination, whereas filament formation is required for germination after prolonged dormancy

Since *S. cerevisiae* harbors two acetyl-CoA synthetase isoforms, we wanted to understand under which specific conditions Acs1 and its polymerization become critical. In an accompanying study, we monitored global protein dynamics throughout meiosis using quantitative mass spectrometry and observed that Acs1 expression is strongly upregulated at the onset of meiosis, when compared to mitotically dividing cells (Figure 6A) ^31^. Acs2, on the other hand, is highly expressed in mitotically dividing cells, but downregulated during meiosis. Western blotting confirmed that the Acs1 level increased in response to pre-meiotic starvation and increased even further approximately four hours after induction of meiosis (Figure 6B).

**Figure 6.**
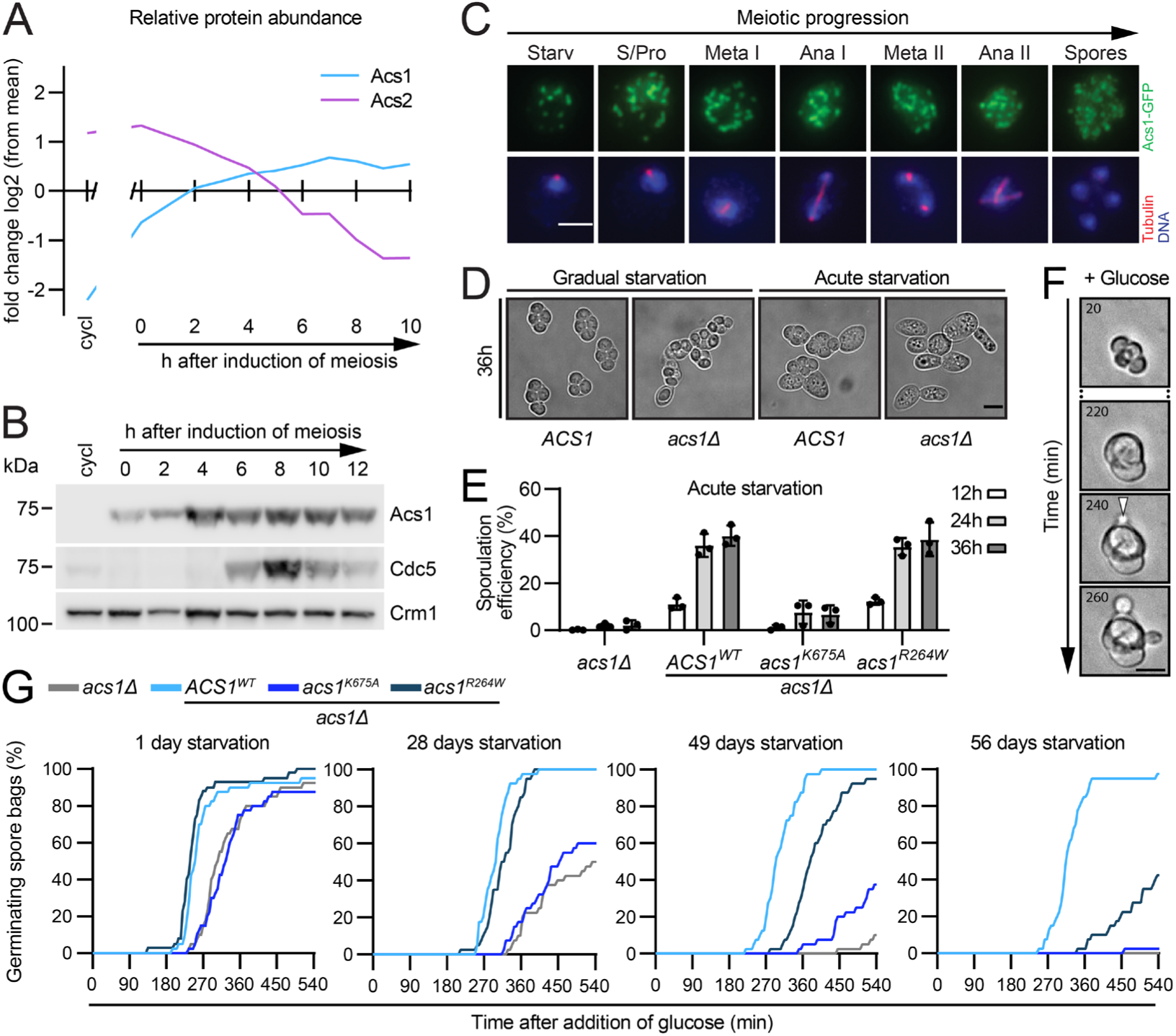
The catalytic activity of Acs1 is required for sporulation and germination, whereas filament formation is required for germination after prolonged dormancy. See also Figure S9-S11. **(A)** Dynamic changes in the relative expression of Acs1 and Acs2 monitored by quantitative proteomics, extracted from ^31^. Samples from cycling mitotic cultures (cycl; in YPD medium), and samples collected at the indicated time points after induction of meiosis from G0/G1 have been analyzed. The plotted values represent the log2 fold change in each sample compared to the average peptide/protein expression from all samples. **(B)** Acs1 is highly expressed throughout meiosis. Western blot analysis of Acs1 in exponentially growing cycling cultures (cycl; in YPD medium), and at the indicated time points after induction of meiosis, in SPM medium. Cdc5 expression was used as a marker of meiotic progression whereas Crm1 serves as protein normalization control. **(C)** Acs1-GFP forms foci and rods throughout gametogenesis. Acs1-GFP expressed from the endogenous promoter was analyzed by immunofluorescence using anti-GFP antibodies. Spindle morphology and DNA were visualized using anti-α-tubulin antibodies and DAPI respectively. The experiment shown is representative for two independent experiments. Scale bar: 3 µm. **(D)** Acs1 is needed for efficient sporulation after acute starvation. Yeast strains with the indicated genotypes were induced to enter meiosis by transfer from YPD to SPM (acute starvation), or by transfer from YPD to YPA to SPM (gradual starvation). Shown are representative images of the sporulation efficiency 36 h after transfer to SPM medium. Note that in acutely starved cell cultures of an *acs1Δ* strain no ascospore formation was observed. Quantification of the sporulation efficiency is shown in Figure S11. Scale bar: 5 µm. **(E)** The catalytic activity of Acs1 is needed for efficient sporulation, whereas filament formation is dispensable. Yeast strains with the indicated genotypes were induced to sporulate as in (**D**) under acute starvation conditions. The efficiency of ascospore formation was quantified by assessing the morphology of 200 cells from three independent experiments after 12, 24, and 36 h in SPM. Ascospore formation was considered successful if at least two spores were enclosed by a spore wall. Plotted values show the mean ± SD. **(F)** Live cell imaging of ascospore germination. Strains shown in (**G**) were induced to sporulate under gradual starvation conditions, as described in (**D**). Mature spores were then induced to germinate by exchanging the sporulation medium (SPM) to glucose-rich medium (SC + 2% Glucose). Light microscopy DIC images were collected every 5 min to follow the initial bud formation, which was considered as the initial point of re-entry into the cell cycle. Example images depict the key morphological changes that occur after transfer of the spores to glucose-rich medium, with bud formation marked by the white arrowhead. Scale bar: 5 µm. **(G)** Acs1 filament formation is needed after prolonged dormancy. Ascospore germination was analyzed as described in (**F**) for strains with the indicated genotypes, after cells were cultured for 1, 28, 49, and 56 days in SPM medium. 40 ascospores per genotype and condition were imaged for cell cycle re-entry. Live cell imaging was performed for a total duration of 540 min (9 h) after glucose addition.

In order to follow the formation of large Acs1 assemblies throughout the meiotic cell division program, we monitored Acs1-GFP by fluorescent light microscopy. Meiotic cells, as well as spores, exhibited fluorescent Acs1-GFP foci or rod-shaped signals (Figure 6C, S9). The number and intensity of Acs1-GFP foci and rods appeared to increase throughout the meiotic program. Correlated light and electron microscopy showed that Acs1-GFP foci indeed colocalized with polymerized Acs1-GFP, even though filament length and filament bundling pattern seemed to be affected by the tag (Figure S10A-C).

Based on the presence of Acs1-GFP foci in meiotic cells as well as in spores, we then probed the role of Acs1 in gametogenesis. We quantified sporulation efficiency in wild-type and different *acs1* mutant strains upon gradual starvation (proliferating cells were starved in YPA before switching to SPM) or acute starvation (proliferating cells were transferred from YPD directly into SPM). No significant differences between the strains tested were observed upon gradual starvation (Figure 6D and S11). In contrast, upon acute starvation, *acs1Δ* cells failed to sporulate (Figure 6E). This was also the case for the catalytic point mutant *acs1^K^*^675^*^A^*. Interestingly, the filament interface mutant *acs1^R^*^264^*^W^*sporulated with comparable efficiency to the wild-type control *ACS1^WT^*.

The experiments above indicate that the catalytic activity of Acs1 is required for efficient gametogenesis, while filament formation is dispensable. However, they do not exclude that filament formation might be required for spore viability, especially after prolonged periods of dormancy. We therefore used live cell imaging to monitor spore germination and return to growth of spores of different ages. Gradual starvation was used to induce sporulation and spores were kept in sporulation medium up to eight weeks before inducing germination by switching to medium containing 2% glucose (Figure 6F). One day after spore formation, most spores from all genotypes germinated within the 9 h of live cell imaging (Figure 6G). However, we noticed a delay in cell cycle re-entry for the *acs1^K^*^675^*^A^* and *acs1Δ* mutants, when compared to the *ACS1^WT^* control. Interestingly, the *acs1^R^*^264^*^W^* spores germinated slightly faster than *ACS1^WT^*, indicating that filament formation might delay the germination of newly generated spores. When spores were kept in sporulation medium for longer periods of time, we noticed a strong reduction in the number of spores from *acs1^K^*^675^*^A^* and *acs1Δ* mutants strains that were able to germinate (Figure 6G). Remarkably, the filament interface mutant strain (*acs1^R^*^264^*^W^*) also showed a steeper increase of time required for re-entry into mitotic proliferation, when compared to the *ACS1^WT^* control. Moreover, after prolonged times of dormancy, a large proportion of *acs1^R^*^264^*^W^* spores completely failed to germinate, whereas the vast majority of *ACS1^WT^* spores re-entered proliferation efficiently (Figure 6G).

These data suggest that the polymerization of Acs1 into filaments is not required for the formation of spores upon starvation. Moreover, filament formation, which inhibits catalytic function, may delay cell cycle re-entry after short periods of starvation. However, failure to form Acs1 filaments results in impaired return to growth after prolonged periods or dormancy. As such, it is likely that the inheritance of meiotically-assembled Acs1 filaments plays a fundamental role in preparing spores for – unpredictably – extended periods of dormancy.

### Acs1 polymerization is a general response to starvation and required for efficient recovery

While monitoring the subcellular localization of Acs1-GFP, we noted that starved pre-meiotic cells already contained Acs1-GFP foci (Figure 6C), suggesting that filament formation may occur as part of a general response to the metabolic state, rather than being a meiosis-specific event. This would be consistent with previous work showing that various GFP-tagged metabolic enzymes, including Acs1, formed fluorescent foci upon nutrient depletion ^10,12^. To test this hypothesis, we cultured haploid cells, which cannot undergo meiosis, for 24 hours in YPA medium (containing acetate, a non-fermentable carbon source, instead of glucose). CryoET imaging of spread spheroplasts revealed prominent Acs1 filament bundles as well as single Ald4 filaments that resembled those observed during meiosis (Figure 7A).

**Figure 7.**
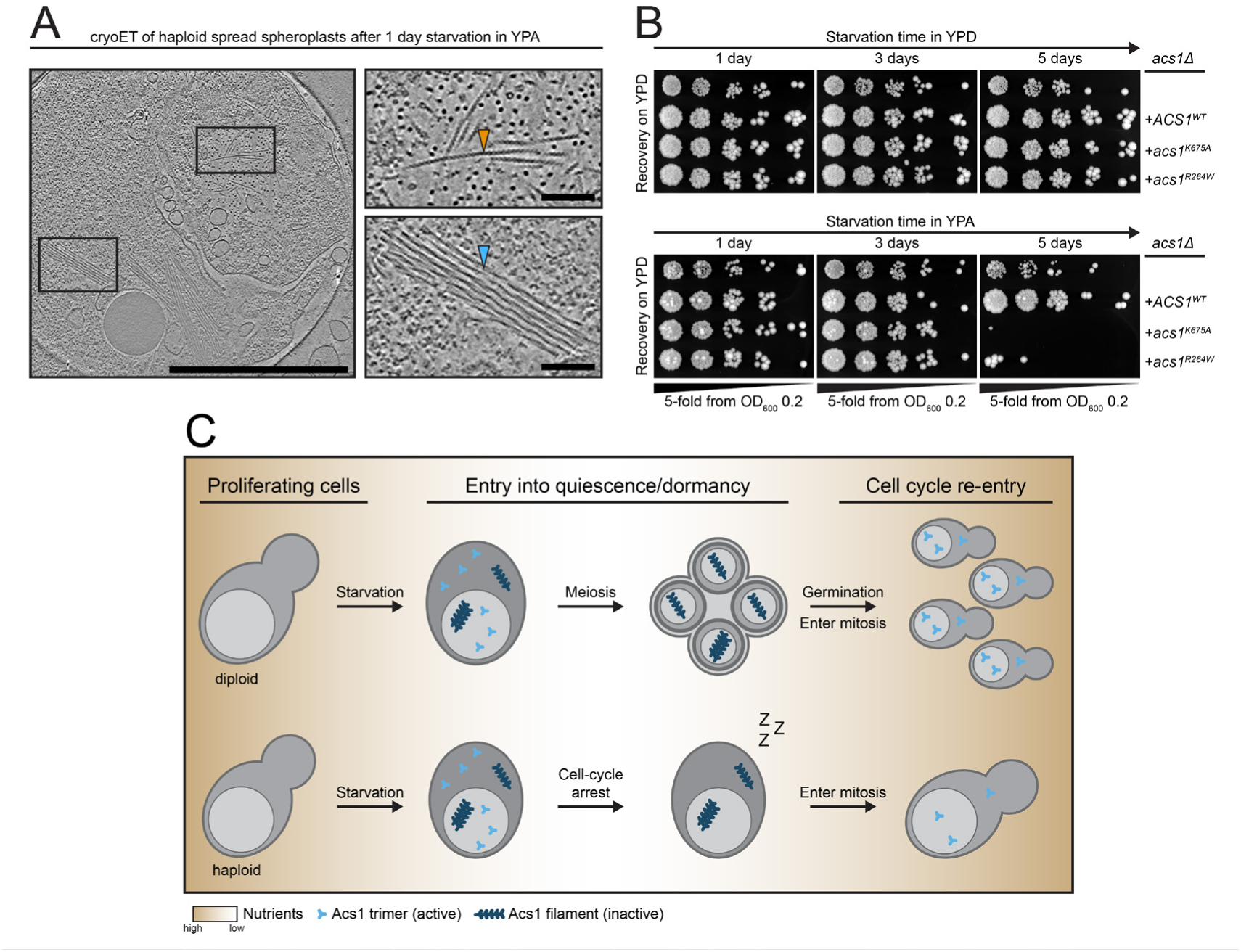
Acs1 polymerization is a general response to starvation and required for efficient recovery. **(A)** Acs1 and Ald4 form filaments also upon starvation independently from meiosis. Haploid yeast cells, which are unable to enter meiosis, were starved for 1 day in YPA, spread onto EM grids, and processed for cryoET. Shown is a 9.02 nm thick slice through a cryo-tomogram with magnified views on the right. Orange arrowhead points to single Ald4 filament whereas the blue arrowhead points to an Acs1 filament bundle. Scale bars: 1 µm and 100 nm for magnified views. **(B)** Acs1 filaments are needed for efficient recovery after starvation. Shown is a cell spotting assay for the indicated genotypes. Cells were incubated for 1, 3, or 5 days in YPD or YPA medium before fivefold serial dilutions were spotted on YPD plates. Note that after five days in YPA medium, deletion strains complemented with *acs1^K^*^675^*^A^* and *acs1^R^*^264^*^W^* almost completely failed to recover. The experiment shown is representative for two independent experiments. **(C)** Model of Acs1 filament function. Starvation triggers entry into quiescence/dormancy via meiosis. In the process, Acs1 expression is upregulated to promote acetyl-CoA production, which is required for meiosis. Shortly after meiotic entry, Acs1 polymerizes and forms filament bundles that are inherited by spores. In the filamentous form, Acs1 is inactive and acetyl-AMP-bound, unable to carry out the second step in acetyl-CoA synthesis. After long periods of dormancy, the stored Acs1 is required for return to proliferation. A similar mechanism operates in haploid cells to enable the return to growth from prolonged periods of starvation.

Having established that Acs1 filament formation also occurs when vegetative cells are starved in medium containing acetate, we tested whether Acs1 played a role in return to growth after starvation, independently from the meiotic cell division program. We cultured haploid cells for different periods of time in medium with glucose (YPD medium) or in medium with acetate (YPA medium). The cells were then transferred onto YPD and return to growth was monitored. As expected, after YPD starvation, wild-type and Acs1 mutants recovered with similar efficiencies (Figure 7B), likely due to the functional compensation by Acs2, which is upregulated in such conditions ^42^. In contrast, after five days of YPA starvation, significant defects in recovery were observed for the Acs1 catalytically deficient mutant (*acs1^K^*^675^*^A^*) as well as for the subunit interface mutant (*acs1^R^*^264^*^W^*). Altogether, these data support a model in which Acs1 filament formation is generally required for the ability of cells to return to proliferation after prolonged periods of starvation.

## DISCUSSION

Our multimodal cryoEM imaging approach on spread samples, termed FilamentID, revealed the structural organization of two novel types of filament assemblies composed of Ald4 and Acs1. Both metabolic enzymes are conserved across all domains of life. Acs homologs play a critical role in carbon metabolism and are also involved in the regulation of histone acetylation, which directly impacts chromatin structure and gene expression ^43,44^. Ald homologs metabolize reactive aldehydes, and, as such, have important roles in detoxification and cell homeostasis ^45^. Reflecting the functions in these central cellular processes, human Acs and Ald homologs are associated with disease. ACSS2 is highly expressed in cancer cells under conditions of metabolic stress ^46^, whereas reduced function of the mitochondrial ALDH2 has been linked to cancer predisposition, cardiovascular disease and, in 8% of the world population, alcohol intolerance ^47^. From this perspective, it will be exciting to investigate whether polymer formation is an evolutionarily conserved aspect of Ald and Acs function, and whether polymerization may become altered in the context of disease. In the following paragraphs, we discuss the impact of our study on the understanding of Acs1 biology and metabolic enzyme filaments in general, as well as the methodological advances.

### Context-specific enzyme polymerization: Acs1 subunit interfaces may sense the metabolic state of cells

Acetyl-CoA synthases and aldehyde dehydrogenases are protein families whose individual isoforms can be differentially regulated in terms of abundance and subcellular localization. Yeast Acs1 and Ald4 share a tight transcriptional repression by glucose ^48,49^. Consistent with this notion, both proteins accumulate in starved pre-meiotic G0/G1 cells and throughout meiosis (Figure 6A) ^31^. Furthermore, among several other metabolic enzymes, Acs1 and Ald4 were previously shown to accumulate as distinct foci in starved yeast cells, when tagged with GFP ^10^. The mechanistic underpinnings of this behavior are currently unknown for most such enzymes. It has been proposed that low nutrient levels lead to cytoplasmic acidification and reduction in cell volume, eventually triggering widespread protein clustering, which leads to the transition of the cytoplasm from a fluid into a solid-like state ^16,50,51^. Acs1 and Ald4 may therefore constitute isoforms of their respective enzyme families that have evolved the ability to polymerize in response to nutrient scarcity.

An elegant example of how a metabolic state can be sensed to trigger filament formation has been proposed for yeast CTP synthase. CTP synthase contains a histidine residue at the subunit interface that becomes protonated at low pH, leading to an increased affinity for the neighboring subunits ^15^. Consistent with the idea that subunit interfaces can sense nutrient levels, our high-resolution structure of natively assembled Acs1 filaments revealed a metabolite, possibly ADP, at the interface of consecutive protomers (Figure 5D). Mutating the binding site at the interface abolished filament bundle formation (Figure 5E, 5G). It is therefore tempting to speculate that sensing of the metabolic state of the cell via ADP levels, which transiently increase during meiosis ^52^, may constitute a trigger for Acs1 to polymerize. This underlines the critical role of subunit interfaces in the response to changes in nutrient availability to tune oligomerization states. Besides this proposed mechanism, it is likely that other regulatory layers play a role, including increased Acs1 expression levels and phosphorylation during meiosis ^31^. These aspects should be considered in future approaches aimed at reconstituting Acs1 filament formation *in vitro*.

### Acs1 polymerization is required for the recovery of chronologically aged dormant cells

We propose a model, in which the essential catalytic function of Acs1, Acetyl-CoA synthesis, is carried out at the onset of meiosis, when ATP production to support the vast biosynthetic needs of sporulation is high ^53^, being shortly after followed by enzyme self-inhibition (Figure 7C). The upregulation of Acs1 under nutrient limiting conditions is accompanied by the polymerization of trimers into filaments, which inactivates the enzyme. Should the inhibition of Acs1 be important for meiosis itself, one would expect that sporulation or spore viability becomes compromised in mutants that specifically fail to form filaments. We found that such a mutant (*acs1^R^*^264^*^W^*) showed normal levels of sporulation (Figure 6E). Remarkably, however, *acs1^R^*^264^*^W^*failed to re-enter the cell cycle after prolonged periods of starvation-induced dormancy (Figure 6G). Therefore, polymer formation appears to sustain the viability of dormant spores that become chronologically old(er). More generally, Acs1 filaments also form in starved haploid cells that cannot undergo meiosis and the ability of Acs1 to polymerize is critical for return to growth after prolonged periods of starvation (Figure 7B).

How does Acs1 polymerization prime dormant/starved cells for efficient return to proliferation? We envision several possible mechanisms: 1) The inactivation of Acs1 in polymers plays a role in lowering acetyl-CoA levels as cells prepare to enter dormancy. This could, for example, ensure that histone acetylation, which is a driver of chromatin openness and gene expression ^54^, remains low. 2) On the other hand, Acs1 depolymerization after prolonged periods of starvation/dormancy, allows for the swift generation of acetyl-CoA by the stored enzyme upon encountering favorable conditions. This provides a two-fold advantage, namely no need for de novo Acs1/Acs2 synthesis, and, since Acs1 filaments are acetyl-AMP-bound, acetyl-CoA can be produced without expending additional cellular ATP. 3) The catalytic inactivation of Acs1, an abundant ATP-consuming enzyme, could help to preserve ATP levels that are needed for minimal metabolism and for the re-entry into a metabolically active state. 4) Storage of Acs1 via the formation of filament bundles could protect from autophagic degradation, which is a highly active process during meiosis ^55^. 5) Acs1 polymerization is part of a general cellular response to solidify the cytoplasm. Such a transition was shown to increase mechanical stability and has been proposed to protect from environmental stress ^50,51^.

An important remaining question is the molecular basis and relevance of Acs1 (and Ald4) to form filaments that are organized in higher-order bundles and arrays (Figure 1A-C). These mesoscale assemblies are likely established through lateral contacts of individual filaments. This idea is supported by the observation that C-terminally GFP-tagged Acs1 formed only short filaments that co-localize in unordered aggregates (Figure S10). The GFP-tag emanating from the filament surface likely disrupts bundle formation and long-range polymerization. These findings underline the importance of integrating cryoET imaging of untagged wild-type cells for a comprehensive understanding of metabolic enzyme filaments across scales. Overall, our study will serve as a framework to understand how the polymerization of metabolic enzymes and the formation of higher-order bundles may prime cells for efficient recovery from dormancy.

### The FilamentID workflow can be adapted to various types of filaments

Besides the possibility of analyzing the *in situ* structure of identified filament assemblies, cryoET is a powerful method to discover new cellular ultrastructures in a range of cellular states, organisms, and tissues^21,22^. The *de novo* identification of such structures *in cellulo*, however, is extremely challenging, as it requires the determination of reconstructions by sub-tomogram averaging at a resolution of typically <4 Å to unambiguously identify unique amino acid sequences. Recent successful examples were enabled by the combination of 1) highly abundant target complexes (e.g. ribosomes) and 2) particularly thin samples for imaging, for instance skinny bacteria ^56,57^ or cryoFIB-thinned cells ^58,59^. An alternative approach to sub-tomogram averaging is to determine high-resolution structures via single-particle cryoEM, however, this is typically done on purified complexes. Integrated with advanced computational analyses, *de novo* identification is also feasible with rather heterogeneous samples ^60–62^. Any purification, however, can lead to disassembly of the structure of interest.

The FilamentID workflow (Figure 2) combines complementary imaging modalities on gently lysed cells/organelles and has three major advantages over the above conventional approaches: 1) FilamentID needs no prior sample knowledge; 2) FilamentID can be applied to targets of rather low abundance and heterogeneity at the superstructural level; and 3) FilamentID sample preparation does not require time-consuming cryoFIB milling or purification optimizations. Limitations of the technique are that 1) the targeted single-particle cryoEM data collection needs to be guided to areas of interest (target complexes need to be visible at low magnification); and 2) membrane integrity and cellular context is lost due to the spreading.

In the present study, FilamentID could unambiguously pinpoint the composition of two different types of filamentous assemblies. We note, however, that we detected other types of filaments in cryo-tomograms of spread yeast cells (Figure S5B). In the future, the workflow could be adapted to study various culturing conditions, distinct developmental programs, different cell types, and diverse organisms.

## ACKNOWLEDGMENTS

We thank the imaging facility ScopeM at ETH Zürich and BioOptics facility at University of Vienna for instrument access and support. The Matos Lab was supported by ETH (31 17-2), SNSF (176108), FWF (SFB Meiosis - 8807-B) and ERC (101002629) grants. The Pilhofer Lab was supported by the NOMIS Foundation.

## AUTHOR CONTRIBUTIONS

J.H. performed the sample preparations for cryoEM/cryoET with the help of F.W.; J.H. and J.X. performed cryoET data collection and analysis; J.H. and J.X. collected cryoEM data; J.X. reconstructed the cryoEM maps and determined the structures; R.W. provided MS data and analysis; J.H., R.W., L.I. D.V. and A.H. generated yeast strains. R.W. and A.H. performed western blotting experiments; J.H. and R.W. performed sporulation efficiency assays; L.I. performed immuno-florescence imaging. J.H. performed correlative cryoLM and cryoET imaging; J.H. did starvation recovery assays; D.V. performed germination assays and analyzed the data with J.H.; J.M. and M.P. conceived and supervised the project; J.H., J.X., J.M. and M.P. wrote the manuscript with input from all authors.

## DATA AND MATERIAL AVAILABILITY

The EMDB entries for the cryo-EM density maps of the Acs1 and Ald4 filaments reported in this paper are EMD-XXXXX (Ald4) and EMD-XXXXX (Acs1). The PDB entries for the corresponding atomic models reported in this paper are XXXX (Ald4) and XXXX (Acs1).

The EMDB entries for the tomograms showing Ald4 or Acs1 filament bundles are EMD-XXXXX (Ald4 filments in mitochondria; meiotic *in cellulo*), EMD-XXXXX (Acs1 filaments in the nucleus; meiotic *in cellulo*), EMD-XXXXX (Acs1 filaments in the cytoplasm; meiotic *in cellulo*), EMD-XXXXX (Ald4 filaments in purified mitochondria from meiotic cells), (Acs1 filaments in spread spheroplasts from meiotic cells), and EMD-XXXXX (Acs1 and Ald4 filaments in spread spheroplasts from starved cells).

## STAR METHODS

### Experimental model and subject details

All strains used in this study are SK1 derivatives and the detailed genotypes are described in Table S2. The following alleles have been described previously: *ndt80Δ, Zip1-(700)-GFP* ^63,64^. Gene deletions were introduced by PCR-based amplification of cassettes from the yeast knock-out collection ^65^. Synthetic *ALD4^WT^* or *ald4^int^*(Lys60Glu, Lys249Glu, Ile250Ala, Gln369Ala, Asn373Ala, Lys377Ala, Asp380Ala, Asn384Ala) constructs including 500 base pairs (bp) upstream of ATG and 500 bp downstream of the terminator were cloned into the plasmid Yiplac128 (pML764) using HindIII and EcoRI restriction sites. The resulting plasmids (pML767 and pML776) were used to reconstitute an *ald4Δ* strain by the integration of the respective linearized vector variant into the promotor region of *ALD4* using SmaI. In all strains carrying *ACS1* variants, a synthetic *ACS1^WT^-6xHIS-6xFLAG* gene sequence construct including 301 bp upstream of ATG and 81 bp downstream of the terminator was cloned into the plasmid pYIplac211 (pML533) using HindIII and EcoRI restriction sites. Plasmids carrying *acs1^K^*^675^*^A^* (pML624) or *acs1^R^*^264^*^W^* (pML619) were generated by site-directed mutagenesis of pYIplac211-*ACS1-6xHIS-6xFLAG* (pML606). Reconstitution of *acs1Δ* strains with *ACS1^WT^* or with the *acs1* point mutants were performed by integration of the respective linearized vector variants into the promotor region of *ACS1* using Bsu36I. For C-terminal PCR-based tagging of chromosomal genes or tag exchange with yeast enhanced GFP (referred to in the text as GFP) cassette was amplified from a plasmid (pML100, Gift from Wolfgang Zachariae) with 50 bp overhangs homologous to 50 bp up- and downstream of the STOP codon ^66^. To remove C-terminal tags, a cassette from pML702 with 50 bp overhangs at the C-terminus including the STOP codon was used. DNA cassettes were transformed into SK1 yeast strains using standard protocols.

### Method details

#### Generation of diploid strains and meiotic time courses

Meiotic time courses were performed as previously described ^23,24^. In brief, haploid cells with opposite mating types were mated on YPD [1% (w/v) yeast extract, 2% (w/v) peptone, 2% (w/v) dextrose]. Diploid cell colonies were then selected on YPG [1% (w/v) yeast extract, 2% (w/v) peptone, 2% (v/v) glycerol] plates for 48 h at 30 °C. Several colonies were picked and expanded on YPD plates for two times 24 h at 30 °C to form a lawn covering the whole plate. Afterwards, cells were inoculated in pre-sporulation medium [YPA, 1% (w/v) yeast extract, 2% (w/v) peptone, 2% (w/v) potassium acetate] at OD_600_ ∼0.3 and grown for 15 h at 25 °C or for 11 h at 30 °C, washed once in sporulation medium [SPM, 2% (w/v) potassium acetate] and inoculated at OD_600_ ∼3.5 into SPM. This time was defined as t = 0 h for all meiotic time courses. Cells were then sporulated at 30 °C and samples were taken at specific time points post transfer to SPM.

#### Sporulation efficiency

To assess sporulation efficiency, cells were inoculated in YPA to OD_600_ ∼0.3 and grown for 15 h at 25°C, washed once in SPM, and inoculated into SPM to OD_600_ ∼3.5. For the acute starvation procedure, cells were directly inoculated into SPM after expansion on YPD plates. An aliquot of cell suspension was collected after 12 h, 24 h, and 36 h in SPM and stored at -20 °C prior to light microscopy imaging. Samples were imaged with standard light microscopy systems. Areas with ascospores and cells were randomly picked, and Z-stacks were recorded in the Brightfield channel. Ascospores and cells were manually scored using Fiji ^67^. 100-200 ascospores or cells were counted in three independent experiments. Ascospore formation was considered successful if the ascus contained at least two spores enclosed by a separate spore wall. Data analysis was performed in Excel (Microsoft) and Prism (version 9.2.0 for Windows; GraphPad) software.

#### Yeast spore germination imaging

Sporulation cultures generated as described above were incubated for up to 56 days at 30 °C in SPM medium supplemented with Ampicillin (50 µg/ml). Every three days, fresh Ampicillin was added. For life cell imaging, samples were adjusted to an OD_600_ ∼1.7 in conditioned SPM before they were put in Concanavalin A (in 1M MnCl_2_, 1x PBS) coated imaging chambers [Nunc™ Lab-Tek™ II Chambered Coverglass, 8 Well, 1.5 Borosilicate Glass (LifeTech Cat 155409)]. After 3 min of incubation at RT, chambers were washed with conditioned SPM before synthetic medium [SC, 0.17% (w/v) yeast nitrogen base w/o ammonium sulphate and amino acids, 0.5% (w/v) ammonium sulphate, 1.1% casamino acids, 0.0055% (w/v) for adenine, tyrosine, uracil, tryptophane and leucin in H_2_O] w/o glucose was added. Right before imaging, SC w/o glucose was exchanged with SC containing 2% (w/v) glucose. Live-cell imaging was performed on a DeltaVision Ultra Epifluorescence Microscope equipped with an Olympus UPlanXApo 40x/0.95 Corr objective, a sCMOS camara (pco.edge 4.2) and a full environmental chamber at 30 °C. 8 µm z-stacks in the DIC channel were collected every 5 min for a total duration of 9 h on pre-selected regions. Additionally, a UV blocking filter was applied.

Movies were analysed with Fiji imaging software. Single ascospores containing at least three visible spores enclosed by a separate spore wall were tracked over the entire movie. The time of initial cell cycle re-entry was defined as the appearance of the first budding spore of an ascus. 40 ascospores per genotype were inspected for each time point. Germination curves were generated in Excel (Microsoft) and Prism (version 9.2.0 for Windows; GraphPad) software.

#### Yeast spot assays

Haploid yeast cells were cultured in liquid YPD over night at 30 °C. In the morning, cells were diluted and inoculated at OD_600_ ∼0.1 in YPD or OD_600_ ∼0.2 in YPA medium supplemented with Ampicillin (50 µg/ml) and incubated at 30 °C. Every three days, cell cultures were supplemented with fresh Ampicillin. For the starvation recovery spot assay, cells were harvested for each time point, washed once in H_2_O and spotted on YPD plates as 5-fold serial dilutions starting from OD_600_ = 0.2. YPD plates were incubated for 48 h at 30 °C before they were imaged with a ChemiDoc Imaging System (BioRad).

#### Protein analyses by western blotting

Samples were processed as described previously (Matos et al., 2008). In short, cell pellets were disrupted in 10% TCA using glass beads on a FastPrep-24 (MP Biomedicals) running two cycles of 40 s (6.5 m/s). The protein precipitates were resuspended in 2X NuPAGE sample buffer and neutralized with 1M TrisBase at a 2:1 ratio (v/v), boiled at 95°C for 10 min and cleared by centrifugation for another 10 min. The relative protein concentration was assessed using the Bio-Rad protein assay. Protein samples were separated on NuPAGE 3-8% Tris-Acetate gels (Invitrogen) and transferred onto Amersham Hybond 0.45 µm PVDF membranes. For immunoblotting the following antibodies were used: mouse anti-GFP (1:2000, 11814460001 Roche), mouse anti-Cdc5 (1:2500, clone 4F10 MM-0192-1-100 MédiMabs), rabbit anti-Acs1 (1:5000, this publication), rabbit anti-Crm1 (1:5000, Onischenko E. et al., 2009). The following secondary HRP conjugated antibodies were used: 1:5000 goat anti-mouse immunoglobulin (P0447 Agilent) and 1:5000 swine anti-rabbit immunoglobulin (P0399 Agilent).

#### FACS analysis of DNA content

Cellular DNA content was determined as described previously ^68^. 1 ml of meiotic cell culture was fixed in 70% (v/v) ice cold EtOH before cells were washed in 50 mM Tris-HCl pH 7.5 and resuspended in 500 µl 50 mM Tris-HCl pH 7.5. RNA was digested o/n or for at least 4 h at 37 °C by adding 2 µl RNase (100 mg/mL) (Roche). Cells were washed in FACS buffer (200 mM Tris-HCl pH 7.5, 211 mM NaCl, 78 mM MgCl_2_), resuspended in 500 µl FACS buffer containing 50 µg/ml propidium iodide and briefly sonicated. An aliquot of 40-60 µl was diluted in 1 ml of 50 mM Tris-HCl pH 7.5. Propidium iodide-stained DNA content was measured on a flow cytometer (FACSCalibur or LSRFortessa, Becton Dickinson). The cytometer data was analyzed using FlowJo software (Becton Dickinson).

#### Fluorescence light microscopy

Immunofluorescence stainings were performed as previously described ^69,70^. The following primary antibodies were used: rabbit anti-GFP (1:600, a gift from Wolfgang Zachariae), rat anti-α-tubulin (1:600, Biorad MCA78G). Secondary antibodies coupled to Alexa 555 and Alexa 488 were used for detection (1:300, Invitrogen). DNA was stained with 4’,6-diamidino-2-phenylindole (DAPI). Images were acquired on a Zeiss Axio Imager M2 equipped with a 63x 1.4 Oil DIC M27 objective and a CoolSnapHQ2 camera under the control of ZEN blue 3.3. Image analysis was performed using Fiji.

For direct fluorescent microscopy without prior staining, ∼8 OD_600_ units were harvested, concentrated, and fixed in 100 µl 4% PFA solution for 20 min at RT without shaking. Samples were than washed in 1x PBS and resuspended for imaging. Images were acquired on a Zeiss Axio Imager M2 equipped with a 63x 1.4 Oil DIC M27 objective and a CoolSnapHQ2 camera under the control of ZEN blue 3.3, or on a Leica Thunder Imager 3D cell culture microscope equipped with a HC PL APO 100x/1.44 Oil CORR CS objective and a sCMOS camara (Leica DFC9000 GTC) under the control of LAS X (v 3.7.6) software. Images acquired at the Leica microscope were processed with large volume computational clearing (LVCC).

#### Isolation of mitochondria and spreading on EM grids

Crude mitochondrial fractions were isolated at a small scale as described previously ^32,71^. Briefly, ∼350 OD_600_ units were harvested, washed in water, and treated with 10 mM DTT in 100 mM Tris-SO_4_ pH 9.4 for 20 min at 30°C. After washing in zymolyase buffer (1.2 M sorbitol, 20 mM KP_i_ pH 7.4), cells were resuspended in 1.5 ml zymolyase buffer containing 10 mg 20T zymolyase (Seikagaku Biobusiness) and incubated for 30 min at 30 °C. Spheroplasts were washed in zymolyase buffer and homogenized in homogenization buffer [0.6 M sorbitol, 10 mM Tris/HCl pH 7.4, 1 mM ethylenediaminetetraacetic acid (EDTA), 2 mM phenylmethylsulfonyl fluoride (PMSF), 0.2% (w/v) bovine serum albumin] by passing the cell suspension 20 times through a 0.8 × 22 mm cannula. Cell debris and nuclei were removed by centrifuging the suspension at 1,000 × g and crude mitochondria were isolated by centrifuging the supernatant at 12,000 × g for 15 min. The mitochondrial pellet was resuspended in SEM buffer (250 mM sucrose, 1 mM EDTA, 10 mM MOPS/KOH pH 7.2) and the protein concentration was estimated against an Albumin standard by the Bradford method. Aliquots were snap-frozen in liquid nitrogen and stored at -80°C or directly used to apply on EM grids for plunge-freezing. For single-particle EM data acquisition, purified mitochondria were splashed by hypo-osmotic swelling. Samples were spun down at 20,000 × g for 10 min and pellets were resuspended in 10 mM MOPS pH 7.2. After 10 min on ice, the mitochondria suspension was plunge-frozen on EM grids.

#### Preparation of spread yeast spheroplasts on EM grids

The preparation of spread yeast spheroplasts on EM grids was adapted from chromosome surface spread protocols ^27,28^. Briefly, ∼3 OD_600_ units of cell culture were harvested, spun down for 2 min at 800 ×g, resuspended in 200 µl of 1.2 M sorbitol in SPM and placed on ice. Cells were spheroplasted by adding 2 µl of 1 M DTT (30 °C, 15 min), followed by 5 µl 100T Zymolyase (1 mg/ml) (Seikagaku Biobusiness) (30 °C, 15-30 min). From 8 min onwards, cells were regularly checked under the microscope to assess the level of spheroplasting by mixing 1 µl of spheroplast suspension with 5 µl water on a glass slide.

Spheroplasted cells appear dark, whereas non-spheroplasted cells have a bright halo. When most of the cell population is spheroplasted, 1 ml of ice-cold STOP solution (0.1 M MES, 1mM EDTA, 0.5 mM MgCl_2_, pH 6.4) was added. Spheroplasts were spun down at 800 × g for 2 min at 4 °C and carefully resuspended in 100-200 µl ice cold STOP solution. 10 µl of spheroplast suspension was then mixed with 40 µl of 1% (v/v) Lipsol or 1% (v/v) NP-40 detergent immediately before adding them onto EM grids for plunge-freezing.

#### Plunge-freezing

Meiotic yeast cells were cultured and plunge-frozen as described previously ^25^ with minor modifications. Meiotic yeast cells were harvested and diluted to an OD_600_ of 1-3 in SPM. Vegetatively growing cells in YPD were harvested at an OD_600_ of ∼0.8 and concentrated in SPM to OD_600_ of 1-3 by spinning for 2 min at 650 × g. Yeast cells were kept on ice until they were plunge-frozen with a Vitrobot Mark IV (Thermo Fisher Scientific) ^72^. 4 µl of cell suspension was applied onto the negatively glow-discharged EM grids (R2/2 Cu 200 mesh, specially treated, Quantifoil). Using a Teflon sheet on one side, grids were back-blotted either once for 5-6 s or twice for 3-5 s at 4 °C, 95% humidity, before plunging into the liquid ethane-propane mixture [37% (v/v) ethane] ^73^. Intact and splashed mitochondria were prepared as described above. Samples were mixed with 10 nm BSA-coated colloidal gold particles (Cytodiagnostics) in a ratio of 5:1 and then 3 µl of sample was applied onto negatively glow-discharged EM grids (R2/2 or R2/1 Cu 200 mesh, specially treated, Quantifoil). Grids were back-blotted for 4-5 sec at 4 °C, 95% humidity and were plunge-frozen. Spread yeast spheroplasts were prepared as described above. The spheroplast Lipsol mixture was mixed with gold particles in a 10:1 ratio and then 3 µl of sample was applied onto the negatively glow-discharged EM grids (R2/1 Cu 200 mesh, specially treated, Quantifoil). Grids were back-blotted for 4-5 sec at 4 °C, 95% humidity and were plunge-frozen.

#### Cryo-focused ion beam milling

Due to the thickness of frozen-hydrated yeast cells, plunge-frozen cells were cryo-focused ion beam (FIB) milled with a Crossbeam 550 FIB-SEM instrument (Carl Zeiss Microscopy) before cryoET imaging as described previously ^25^. The FIB-SEM instrument was equipped with an SE2 detector (Carl Zeiss Microscopy), an in-lens secondary electron detector (Carl Zeiss Microscopy), a copper band-cooled mechanical cryo-stage (Carl Zeiss Microscopy), and an integrated VCT500 vacuum transfer system (Leica Microsystems). In brief, plunge-frozen EM grids were clipped into FIB milling Autogrids (Thermo Fisher Scientific) and mounted onto a pre-tilted cryo-FIB Autogrid holder (Medeiros et al., 2018) (Leica Microsystems) using a VCM loading station (Leica Microsystems). Using the VCT500 shuttle, the Autogrid holder was transferred to an ACE600 (Leica Microsystems) to cryo-sputter-coat the sample with a 4 nm thick layer of tungsten. Afterwards, the samples were loaded into the Crossbeam 550 using the VCT500 vacuum transfer system. Inside the FIB-SEM, grids were additionally coated with organoplatinum. Targets were selected with the SEM and automated sequential FIB milling was set up. A pattern with four currents was used (rough milling: 700 pA, 300 pA, and 100 pA; polishing: 50 pA) for milling, targeting for a ∼300 nm thick lamella. After milling, the Autogrid holder was transferred back to the VCM loading station with the VCT500 shuttle and grids were unloaded and stored in liquid nitrogen until cryoET imaging.

#### CryoLM

Plunge-frozen EM grids were imaged with a Zeiss LSM900 equipped with Airyscan2 detector and a Linkam CMS196V3 cryo-stage in a de-humidified room. EM grid overviews were collected with a 5x/0.2 DIC C Epiplan-Apochromat objective to localize the regions of interest. Afterwards, z-stacks were collected with a 100x/0.75 DIC LD EC Epiplan-Neofluar (WD = 4 mm) objective. Confocal imaging tracks were used to visualize Acs1-GFP signal as well as the EM grid with transmitted and reflective light. Confocal imaging stacks were deconvolved with Zeiss LSM Plus processing and maximum intensity projections were generated in ZEN Blue (Carl Zeiss Microscopy, v.3.5) software.

CryoLM data was then used to correlate targets for cryoET data collection in x-y dimensions (described below) ^74^. For that, cryo-LM images were imported into SerialEM ^75^ and aligned to the corresponding EM overviews based on prominent landmarks. To visualize correlated cryoLM and cryoEM images after data collection, low-magnification EM overviews were low-pass filtered using ‘mtffilter’ and converted to tiff files using ‘mrc2tif’ in IMOD package ^76^. These images together with the cryoLM data were imported in ZEN Connect (within ZEN Blue, Carl Zeiss Microscopy, v.3.5) for correlation with the Point Alignment Wizard.

#### CryoET data collection, reconstruction, and segmentation

CryoET datasets were collected on Titan Krios transmission electron microscopes (TEM) (Thermo Fisher Scientific) operating at 300 kV and equipped with Quantum LS filter and K2 Summit direct electron detectors (Gatan) (Krios1), equipped with Quantum LS filter and K3 direct electron detectors (Gatan) (Krios2), and BioContinuum imaging filter and K3 direct electron detectors (Gatan) (Krios3). Low magnification overviews were recorded for navigation and targets were selected for the subsequent tilt series collection using SerialEM ^75^.

For lamellae, tilt series collections were performed using a bidirectional scheme with an angular range between ±70° to ±50°, depending on the pre-tilted geometry of lamella, with 2° increment at a defocus of -8 µm. The pixel sizes in different datasets were 4.34 Å (Krios1) or 4.51 Å (Krios3) or 4.57 Å (Krios2) at the specimen levels. The accumulated dose per tilt series is ∼120 e^-^/Å^2^. For purified mitochondria, tilt series collections were performed using a dose-symmetric scheme with an angular range between +60° to -60° with 2° increment at a defocus of -8 µm. The pixel size was 2.75 Å (Krios1) or 2.68 Å (Krios3) at the specimen level and the total dose per tilt series is ∼160 e^-^/Å^2^. For spread yeast spheroplasts, the same collection scheme was used as for purified mitochondria and the pixel sizes at the specimen level were the same as for the lamella. The total dose per tilt series is ∼130 e^-^/Å^2^.

Frames were aligned using ‘alignframes’ and tomograms were reconstructed using the IMOD package^76^. Tomograms shown in the figures were binned at the level of 4 and the contrasts were improved using the deconvolution filter ‘tom_deconv’ ^77^.

AI-based segmentations were generated in Dragonfly software (Object Research Systems, v. 2022.2) as described previously ^78^. Briefly, filtered tomograms were loaded into Dragonfly and further processed by histogram equalization, Gaussian and Unsharp filtering. Afterwards, a U-Net (with 2.5D input of 5 slices) was trained to recognize background voxels, filaments, and membranes within the tomograms. All AI-segmentations were manually cleaned up in Dragonfly, exported as binary tiff files and converted to mrc files using ‘tif2mrc’ in IMOD. Segmentations were visualized in UCSF ChimeraX ^79^.

#### Sub-tomogram averaging

Sub-tomogram averaging of filaments from different datasets was performed in Dynamo software ^80^. The individual filaments were manually picked from reconstructed tomograms at a binning factor of 4 using the ‘filamentWithTorsion’ model in Dynamo. The different filaments were segmented with the corresponding inter-segment distance (8.8 nm for Ald4 dataset, 5.5 nm for Acs1 dataset) before particle cropping. The cropped sub-volumes from each dataset (Ald4 filaments in the isolated mitochondria: 564 particles from 3 tomograms; Acs1 filaments in the spread yeast spheroplasts: 452 particles from 3 tomograms) were assigned random azimuth orientations using ‘dynamp_table_randomize_azimuth’ and were firstly subjected to the averaging without imposing symmetry. The symmetry information of individual filaments was then deduced from the asymmetric reconstruction and was applied in the downstream averaging analyses. All particles were subjected to one round of coarse alignment with rough angular search steps, and the particles were then split into half-datasets based on the odd-and-even order with ‘dteo’ package in Dynamo. Each half-dataset was subjected to fine alignment with precise angular search steps against the same reference. The Acs1 dataset was analyzed using fine alignment at the binning factor of 2, while the Ald4 dataset was analyzed at the binning factor of 4. The final averaged volumes from individual half-datasets were used to estimate the resolution based on the Fourier shell correlation (FSC) ^81^ using ‘relion_postprocess’ ^82^. The final resolution of averaged maps of Ald4 filaments from intact mitochondria was 44 Å, whereas the resolution of Acs1 filaments from spread yeast spheroplasts was 18.3 Å.

#### CryoEM single-particle data collection

CryoEM data collection parameters are summarized in Table S1. CryoEM datasets of Ald4 filaments from spread yeast mitochondria were collected at a nominal magnification of 81,000 × (an effective pixel size of 0.55 Å at super-resolution) using the SerialEM program on Krios2. Low magnification overviews were recorded for navigation purposes to target filaments and the data was collected as movie stack in super-resolution mode, with the 2.5 s total exposure time at a defocus value from -1.2 to -2.8 µm. Each stack contains 50 frames, and the accumulated electron dose was ∼60 e^-^/Å^2^. The frames of the stack were aligned and applied with dose weighting at the binning factor of 2 using ‘MotionCor2’ ^83^ (an effective pixel size of 1.1 Å). The CTF parameters of micrographs were estimated using ‘Gctf’ ^84^. A total of 531 micrographs in two batches were collected for image processing.

CryoEM datasets of Acs1 filaments from spread yeast spheroplasts were collected at a nominal magnification of 130,000 × (an effective pixel size of 0.535 Å at super-resolution) using the SerialEM program on Krios1. The data collection strategy was performed same as above. The data was collected as movie stack in super-resolution mode, with the 8 s total exposure time. Each stack contains 32 frames and the accumulated electron dose was ∼60 e^-^/Å^2^. The motion correction and CTF parameter estimation were performed the same as above (an effective pixel size of 1.07 Å). A total of 1,084 micrographs were collected for image processing.

#### CryoEM image processing, identification of Ald4, and structural modeling

To determine the structure of the filament observed in the mitochondria, 132 micrographs in the first batch of dataset were used to generate an initial model (Figure S3A). Briefly, the filaments were manually picked using Relion 3.0 ^85^ and 22,365 segments were extracted with an inter-box distance of 42 Å. One round of 2D classification at the binning factor of 4 was performed to remove bad particles, and the particles in good classes were then subjected to 3D refinement without imposing helical parameter but with 2-fold symmetry that was deduced from the sub-tomogram averaged volume. The initial helical parameters were interpreted from the reconstructed map using ‘relion_helix_toolbox’ ^86^ and were then applied and locally refined in the following helical reconstruction ^86^. The bad particles were further removed by one round of 2D classification without sampling after helical reconstruction and the following 2D classification. A total of 11,863 particles were used to reconstruct ∼6.8 Å resolution of filament structure at the binning factor of 2, which was imposed a 2-fold symmetry and helical parameters (twist = 96.2°, rise = 42.2 Å).

To identify the components in the filament assembly, the molecular weight of the sought-after protein was firstly estimated to be ∼30-70 kDa based on the volume size of one filament subunit. To shortlist potential candidates, we cross-compared mass spectrometry (MS) data of mitochondrial matrix/inner membrane proteins ^32^ and MS data covering protein expression profiles in meiosis ^31^ (Table S3). Providing this list to structural docking of Alphafold ^29,30^ predicted candidate protein structures showed that the Ald4 protein fits the map.

To further verify the protein identity, we continued image processing using 531 micrographs collected in two batches. Since Ald4 homolog protein was reported to form a tetramer in solution and crystal structure (Aldh2, PDB entry: 1NZZ) ^33^, there were still two symmetry possibilities for the filament assembly: proteins form a dimer (C2 symmetry) or tetramer (D2 symmetry) and then assemble into the filament. The different symmetry possibilities would request different inter-box distance for filament segmentations.

To investigate this, we first assumed C2 symmetry and segmented the filaments with an inter-box distance of 42.2 Å. The extracted segments were subjected to 2 rounds of 2D classifications at the binning factor of 4 and 2, and the particles in good classes were used for helical reconstruction at the binning factor of 1, where the helical parameters (twist = 96.2°, rise = 42.2 Å) were applied and optimized during refinement. A final 4.2 Å resolution filament structure was determined, however, the densities around the secondary structure (e.g., α helices) were unreasonable, suggesting that a wrong symmetry was applied. Thus, we switched to D2 symmetry in the following image processing. The filaments were segmented with an inter-box distance of 84.4 Å and were extracted with a larger box size (320 pixels). The particles were subjected to a round of 2D classification at the binning factor of 4 and the particles in good classes were used for the helical reconstruction at the binning factor of 1. A filament structure with 6.6 Å resolution was determined, assuming C2 and helical symmetry (twist = 96.2°, rise = 84.6 Å). Moreover, 2D classification analysis revealed that tetramers adopt two different types of stacking for the filament assembly: helical and non-helical (Figure S3B). To further improve the resolution, the central tetramer in each segment was centered and re-extracted using a smaller box size (200 pixels) (Figure S3A). The particles with smaller box sizes were than subjected to 3D refinement followed by local 3D classification. The particles in good 3D classes were combined and were subjected to the next round of 3D refinement imposed C2 and local symmetry (D2). One round of CTF parameter optimization and particle polishing were further applied to improve the map quality. A final resolution of 3.8 Å structure was determined from 29,307 particles assuming C2 and local symmetry (together in D2 symmetry). The map quality was further improved using ‘deepEMhancer’ ^87^. The local resolution was estimated using Relion (Figure S3C).

The structure was manually refined in COOT ^88^, the models were further refined in real-space using iterative refinements of RosettaCM ^89^ and ‘phenix.real_space_refine’ ^90^. The refined model was evaluated using ‘phenix.molprobity’ ^90^ and the model vs. map FSC was calculated using ‘phenix.mtrifage’ ^90^ (Figure S3D-E). In the final atomic model of an Ald4 filament subunit, 24 residues of the N-terminus were missing due to invisible densities. The bound NAD molecule was included in the final model (Table S1).

The surface contact sites between Ald4 tetramers were firstly analyzed by PDBePISA [’Protein interfaces, surfaces and assemblies’ service PISA at the European Bioinformatics Institute. (http://www.ebi.ac.uk/pdbe/prot_int/pistart.html), ^91^] and were then manually checked in COOT.

#### CryoEM image processing, identification of Acs1, and structural modeling

To generate the initial filament model and estimate the helical parameters, micrographs of 0° tilt in the datasets of spread yeast spheroplasts were used for the processing (Figure S6A). The filaments were picked manually with the start-end coordinate pairs using Relion 3.0. The filaments were segmented with an inter-box distance of 60 Å and were then subjected to 2 rounds of 2D classifications. The particles in the good classes were selected for 3D auto-refinement using helical reconstruction without imposing helical symmetry but with 3-fold symmetry that was deduced from sub-tomogram averaged volume. The helical parameters were estimated from the initial reconstruction in real space using ‘relion_helix_toolbox’ and were optimized for the second round of 3D auto-refinement at the binning factor of 4. The refined helical parameters were: twist = 13.58°, rise = 54.62 Å.

To determine the high-resolution filament structure, 1,084 micrographs were collected as mentioned above and used for the imaging processing. The filaments were manually picked and 108,349 segments were extracted with an inter-box distance of 55 Å. Bad particles were removed through 2 rounds of 2D classifications at the binning factors of 4 and 2 (Figure S6B). The particles in good classes were then applied to the helical reconstruction at the binning factor of 2, where the helical parameters from the reconstruction of 0° tilt were applied and locally refined in the follow-up processing steps. One round of 2D classification without sampling was performed after helical reconstruction to remove the mis-aligned particles. The remaining particles were subjected to the 3D helical reconstruction at the binning factor of 1 and a subsequent round of 2D and 3D classifications without sampling. The particles from good classes were then subjected to another round of 3D classification and only one 3D class (class IV in Figure S6B) was used for the helical reconstruction. One round of CTF parameters optimization and particle polishing were further performed to improve the map quality. The final 3.5 Å resolution filament structure was determined from 17,169 particles imposing 3-fold symmetry and helical parameters (twist = 13.03°, rise = 53.61 Å) (Figure S6B). The map quality was further improved using ‘deepEMhancer’. The local resolution was estimated using Relion (Figure S6C).

To identify the components in the filament assembly, the Cα backbone of one filament subunit was manually traced in COOT and was then subjected to DALI search (http://ekhidna2.biocenter.helsinki.fi/dali/) ^35^, which revealed the Acs1 protein as a potential candidate. The crystal structure of Acs1 protein (PDB entry: 1RY2) ^36^ was then docked into the map and manually refined using COOT and then subjected to the same refinements and structural validation as mentioned above for Ald4 (Figure S6D-E). In the final atomic model of Acs1 filament subunit, 37 residues in the N-terminus were missing due to invisible densities. The intermediary product acetyl-AMP was included in the final model (Table S1).

## SUPPLEMENTARY FIGURES

**Figure S1.**
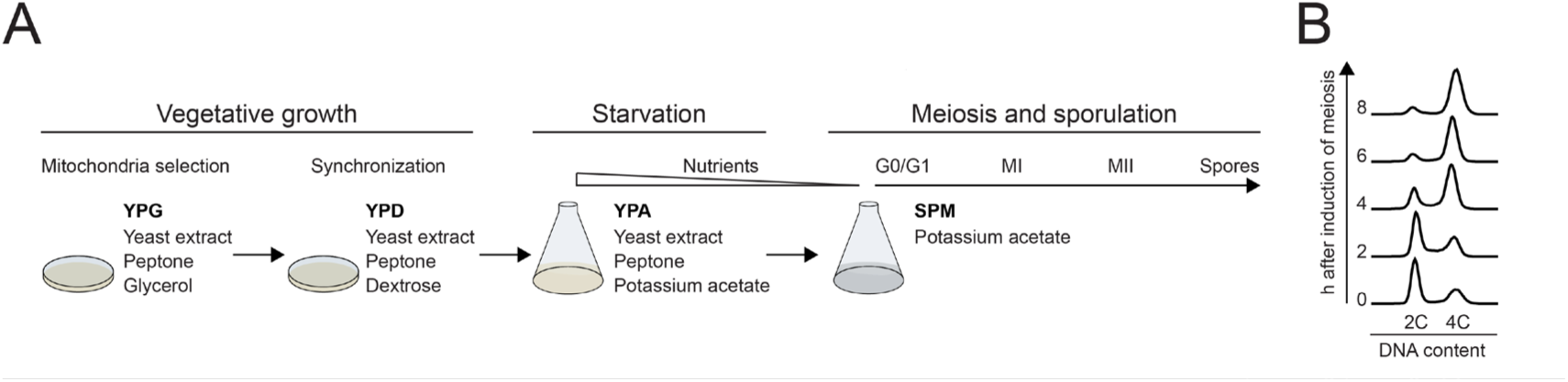
Meiotic time course. **(A)** Schematic representation of the experimental setup to perform a meiotic time course. Vegetatively growing diploid *S. cerevisiae* cells were plated for single colonies on YPG plates to select for strong respiratory growth. Single colonies were then expanded before they were starved in YPA medium containing the non-fermentable carbon source potassium acetate. Arrested cells at G0/G1 were transferred to sporulation medium (SPM) lacking the nitrogen source in order to induce the meiotic cell division program (Pre-meiotic G0/G1, first meiotic division MI; second meiotic division MII, and spore formation). **(B)** FACS analysis of the DNA content to monitor meiotic progression. DNA content of meiotic cell cultures was analyzed by FACS at regular time intervals after the induction of meiosis (transfer of cells to SPM). Note that most of the cells duplicated their genomes after ∼4 h in SPM. Shown is a representative time course.

**Figure S2.**
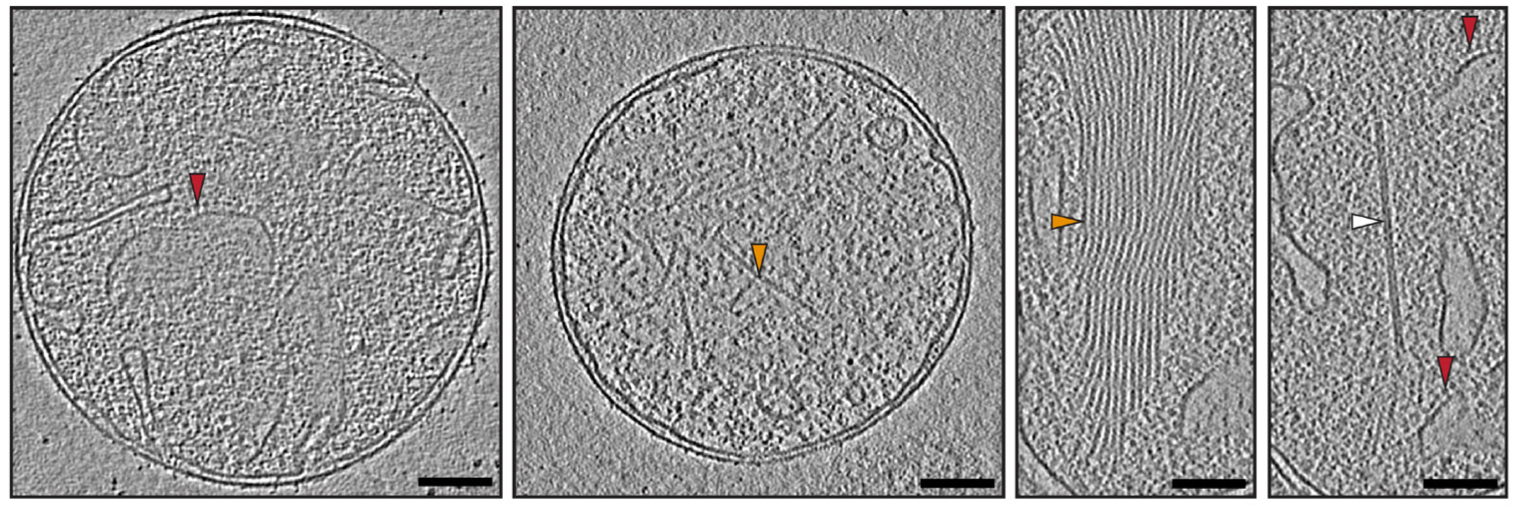
CryoET of purified meiotic mitochondria. Visualization of the ultrastructure of purified mitochondria. Shown are example slices through cryo-tomograms of purified mitochondria from meiotic cell cultures collected between 6-8 h after induction of meiosis. Red arrowheads point to putative F_O_F_1_-ATP synthases on cristae. Orange arrowheads show single filaments and arrays as seen in Figure 3A, whereas the white arrowhead points to a different type of filament within purified mitochondria. Shown are projections of 5.5 nm thick slices. Scale bars: 100 nm.

**Figure S3.**
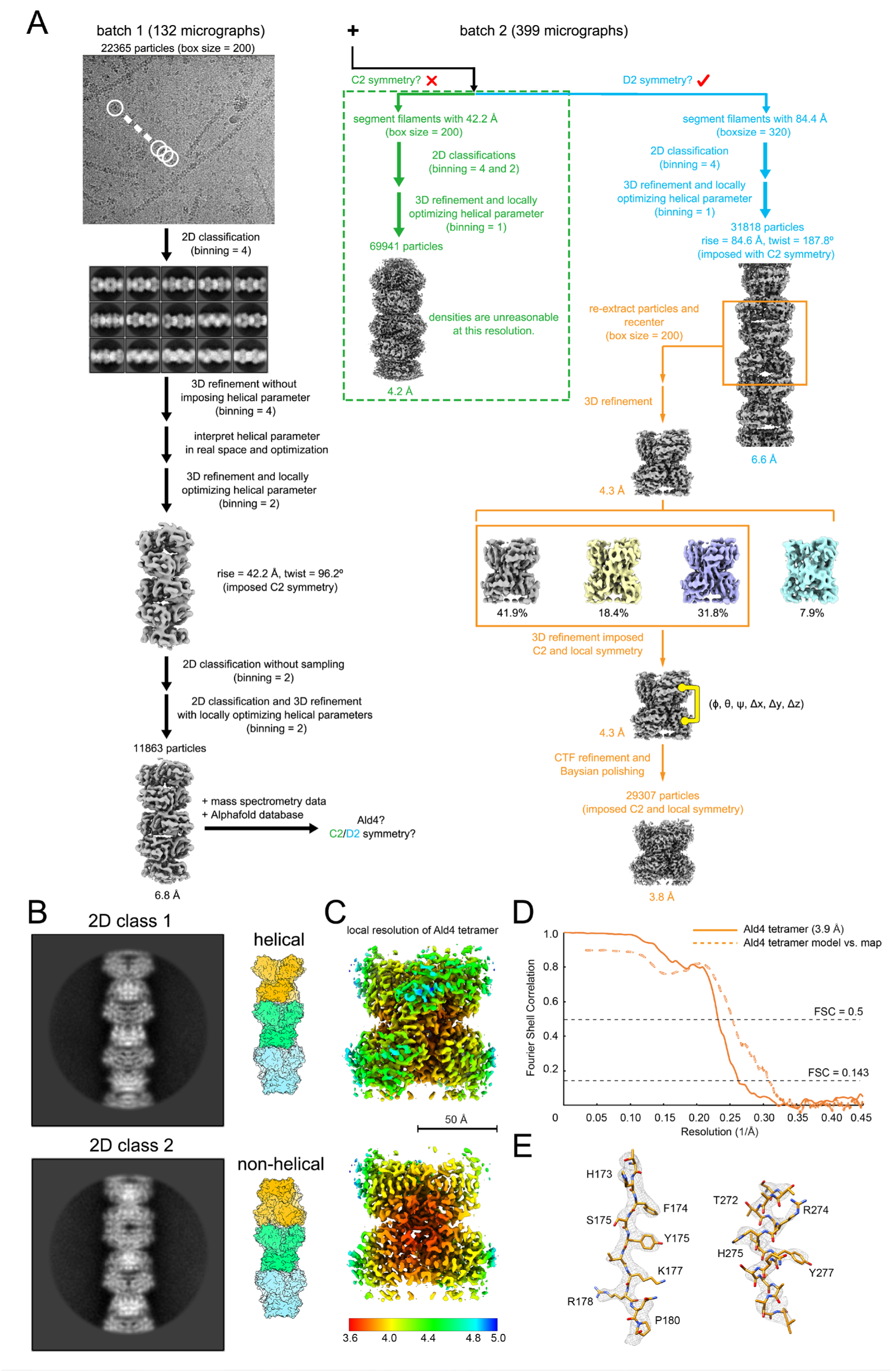
Flowchart of processing cryoEM data for Ald4 filaments. **(A)** Flowchart for the cryoEM reconstruction of the Ald4 filament. See METHODS and Supplementary Table S1 for details. **(B)** Representative 2D classes and the corresponding schematics showing that Ald4 tetramers follow two different stacking manners for the filament assembly: helical (top) and non-helical (bottom). Individual tetramers are colored in orange, green, and blue respectively. **(C)** Local resolution (indicated by colors in Å) maps of the Ald4 tetramer. Scale bar: 50 Å. **(D)** Plots showing the gold standard FSC curve of the cryoEM reconstruction of the Ald4 tetramer (orange) and the corresponding model *vs.* map FSC curve (dashed orange). **(E)** Stick and density mesh diagrams showing representative density maps of the Ald4 tetramer. Stick models and density meshes are colored orange and gray respectively.

**Figure S4.**
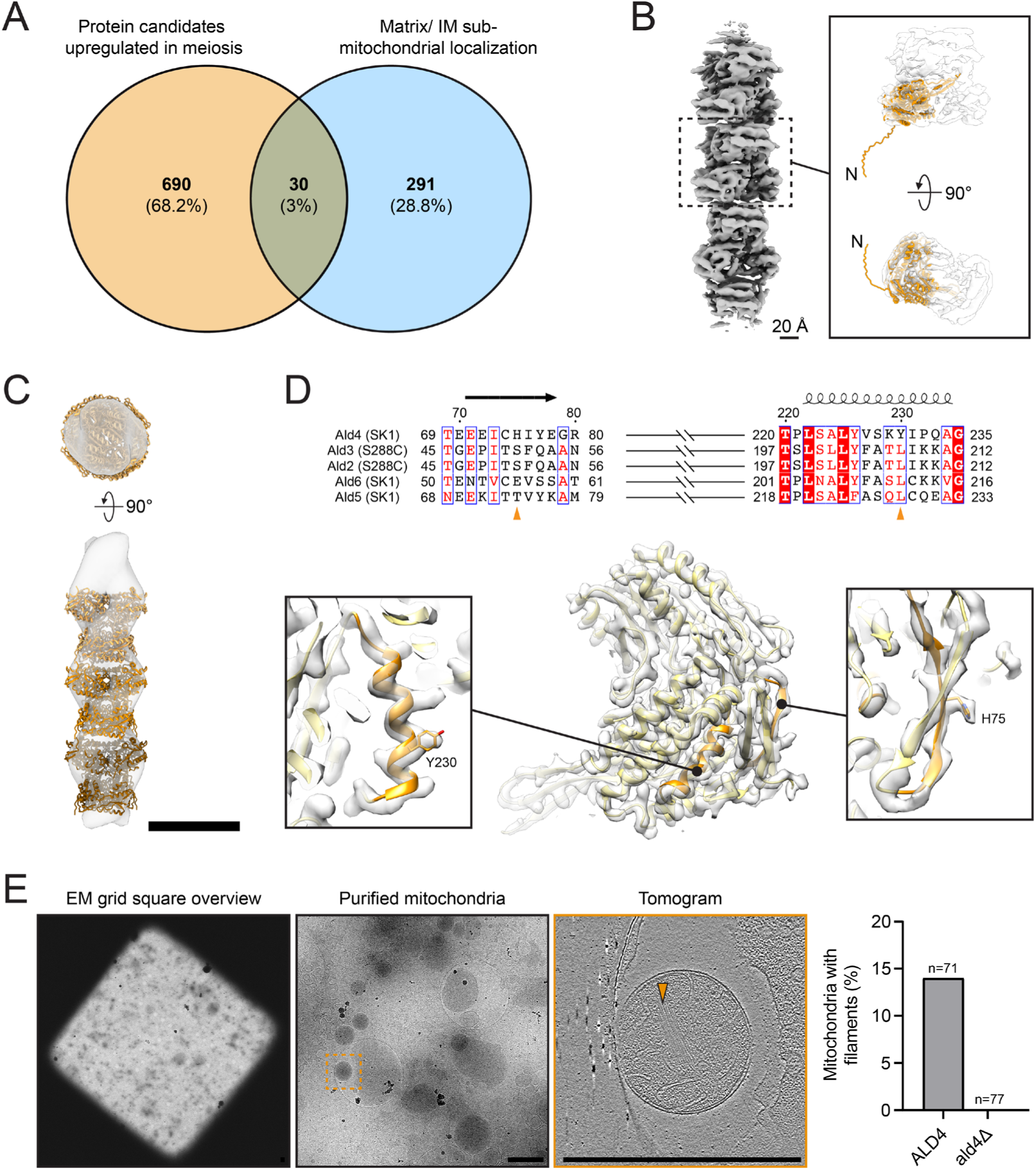
Validation of Ald4 protein identity. **(A)** Comparing data obtained from two independent mass spectrometry analyses yields 30 candidate proteins forming the mitochondrial filaments. Shown is a Venn diagram with proteins upregulated in meiosis ^31^ colored orange together with high confidence mitochondria proteins, which localize to the mitochondrial matrix or inner membrane (IM) colored blue ^32^. Percentage of total input is indicated in each bin. **(B)** Ribbon and shadowed surface diagrams showing that Ald4 is a potential candidate from (**A**) forming the mitochondrial filaments. Left: the helical reconstructed map of the mitochondrial filament. One helical repeat is highlighted with a dashed black box. Right: docking of the predicted Ald4 structure from Alphafold ^29,30^ (orange) into one subunit of one helical repeat. **(C)** Docking of the high-resolution structure of three Ald4 tetramers with the sub-tomogram averaged volume of Ald4 filaments from splashed mitochondria. Three layers of Ald4 tetramers are shown as ribbons and are colored in different shades of orange as in Figure 4D, while the averaged volume (also shown in Figure 3B) is transparent and colored gray. Scale bar: 10 nm. **(D)** Reconstructed map unambiguously identifies Ald4 as the only filament component. Top: sequence alignments of different Ald isoforms (Ald4-6 from SK1 strain, Ald2-3 from S288C strain). Identical residues are shown in white on a red background, while similar residues are shown in red. The blue boxes indicate the conserved positions. The secondary structure of Ald4 is shown above the corresponding sequences. The image is made using Espript ^92^. Two distinguishing residues of Ald4 (H75 and Y230) are highlighted by orange arrowheads and are shown in the bottom panel. Bottom: ribbon and shadowed surface diagrams showing the fitting of the Ald4 model with the reconstructed map. The density map is colored gray and is shown transparent, while the overall model is colored yellow. The aligned parts shown on the top panel are colored orange, while zoom-ins show the densities of two distinguishing residues (H75 and Y230). **(E)** Mitochondrial filament arrays consist of Ald4. Left to right: to check for the presence of filaments, purified mitochondria from meiotic cell cultures are picked in the EM grid square overview followed by cryoET imaging. The corresponding 5.5 nm thick slice of the cryo-tomogram shows an example mitochondrion containing filaments. Scale bars: 1 µm. Right: shown are percentages of meiotic mitochondria (6 h after induction of meiosis) containing filaments and the total number of mitochondria (n) analyzed for each genotype are indicated.

**Figure S5.**
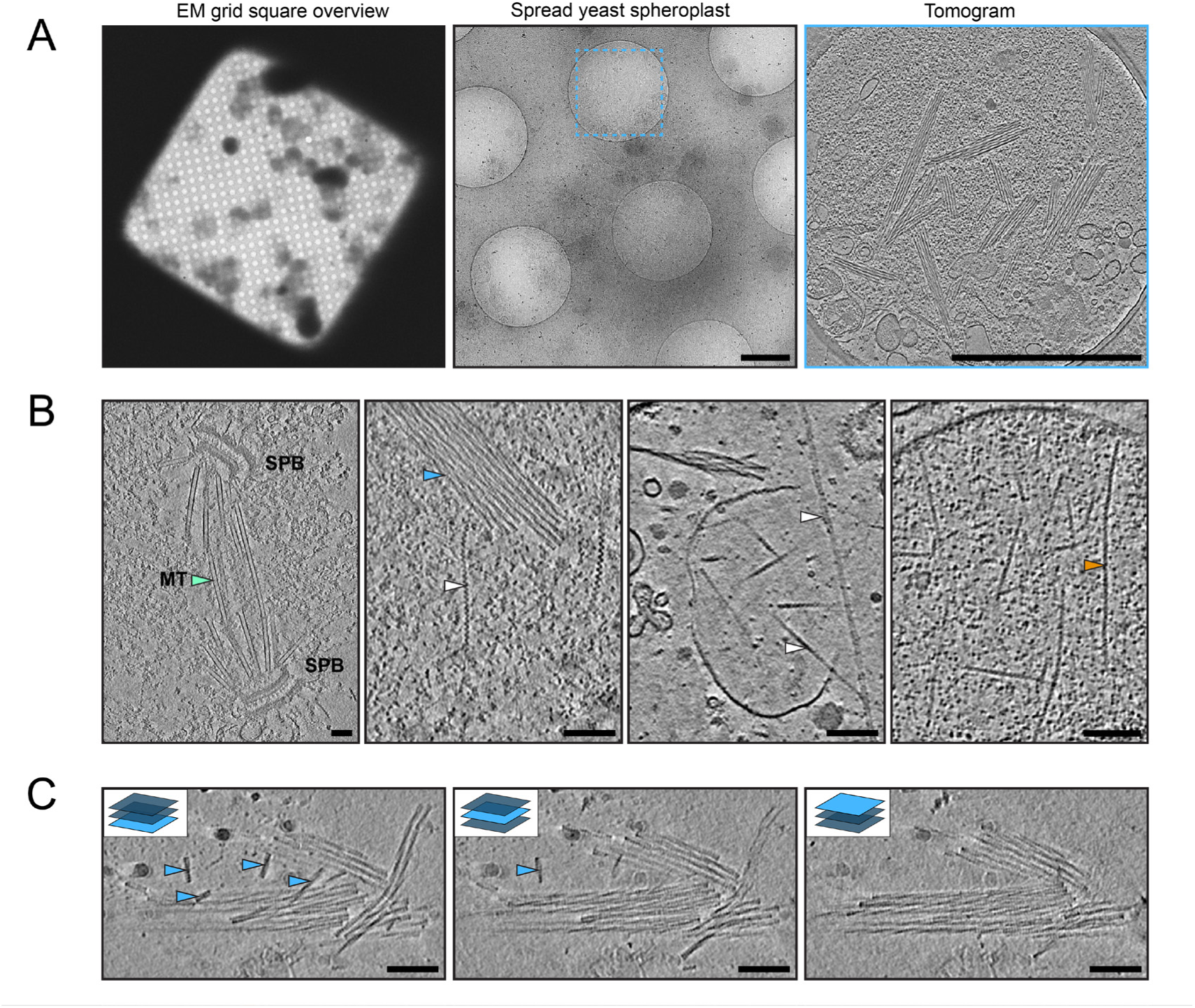
CryoET on spread spheroplasts. **(A)** Spread yeast spheroplasts allow for direct cryoET imaging. Left: overview of an example EM grid square with spheroplasts. Middle: cryoEM micrograph of an example spread spheroplast from a *ndt80Δ* strain arrested in prophase I, 8 h post induction of meiosis. Right: slice through the cryo-tomogram collected on parts of the spread indicated by the blue box. Note that straight filament assemblies in different directions can be observed as shown in Figure 4A. Shown is a projection of 9.14 nm thick slices. Scale bars: 1 µm. **(B)** Spread yeast spheroplasts enable the visualization of various macromolecular assemblies. Shown are example slices through cryo-tomograms of spreads from meiotic cell cultures. Green arrowhead points to microtubules (MT) branching from spindle pole bodies (SPB). Blue arrowhead shows filament bundles, orange arrowhead shows single filaments in spread mitochondria, white arrowheads point to other types of filaments. Shown are projections of either 8.68 nm or 9.14 nm thick slices. Scale bars: 100 nm. **(C)** Slice through cryo-tomogram of potential filament bundle (dis-)assembly intermediates. Note that short single filaments point towards a bundle containing multiple long filaments. Short filaments are highlighted with blue arrowheads. Shown are projections of 9.14 nm thick slices at different z-heights of the tomogram, which are 7.31 nm apart. Scale bar: 100 nm.

**Figure S6.**
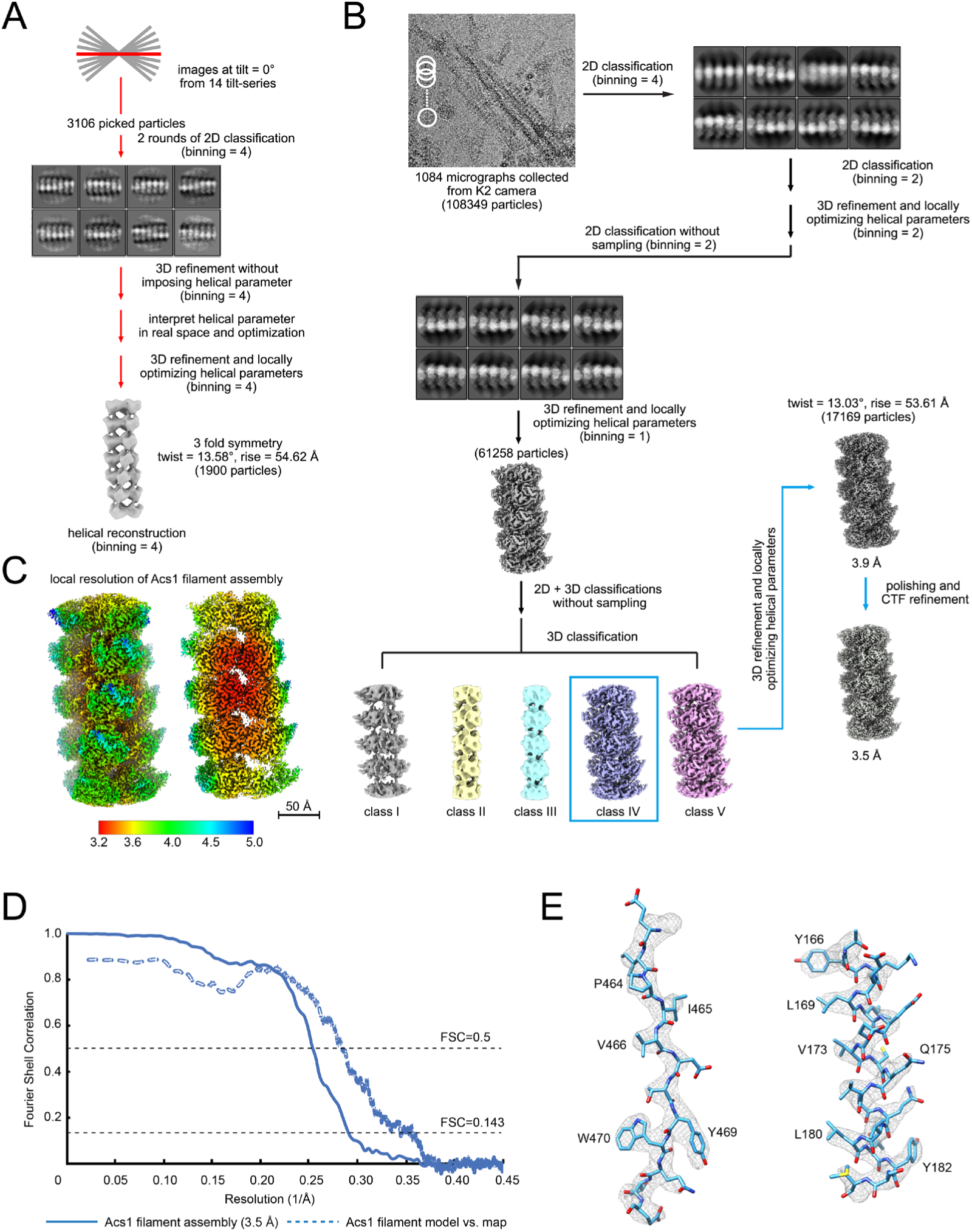
Flowchart of processing cryoEM data for Acs1 filaments. **(A)** Flowchart to generate the initial filament model and to determine the initial helical parameters using individual images (tilt = 0°) from different tilt series. See METHODS for details. **(B)** Flowchart for the cryoEM reconstruction of Acs1 filament. See METHODS and Supplementary Table S1 for details. **(C)** Local resolution (indicated by colors in Å) maps of the Acs1 filament structure. Scale bar: 50 Å. **(D)** Plots showing the gold standard Fourier Shell Correlation (FSC) curve of the cryoEM reconstruction of Acs1 filament (blue) and the corresponding model *vs.* map FSC curve (dashed blue). **(E)** Stick and density mesh diagrams showing the representative density maps of the Acs1 filament structure. Stick models and density meshes are colored blue and gray respectively.

**Figure S7.**
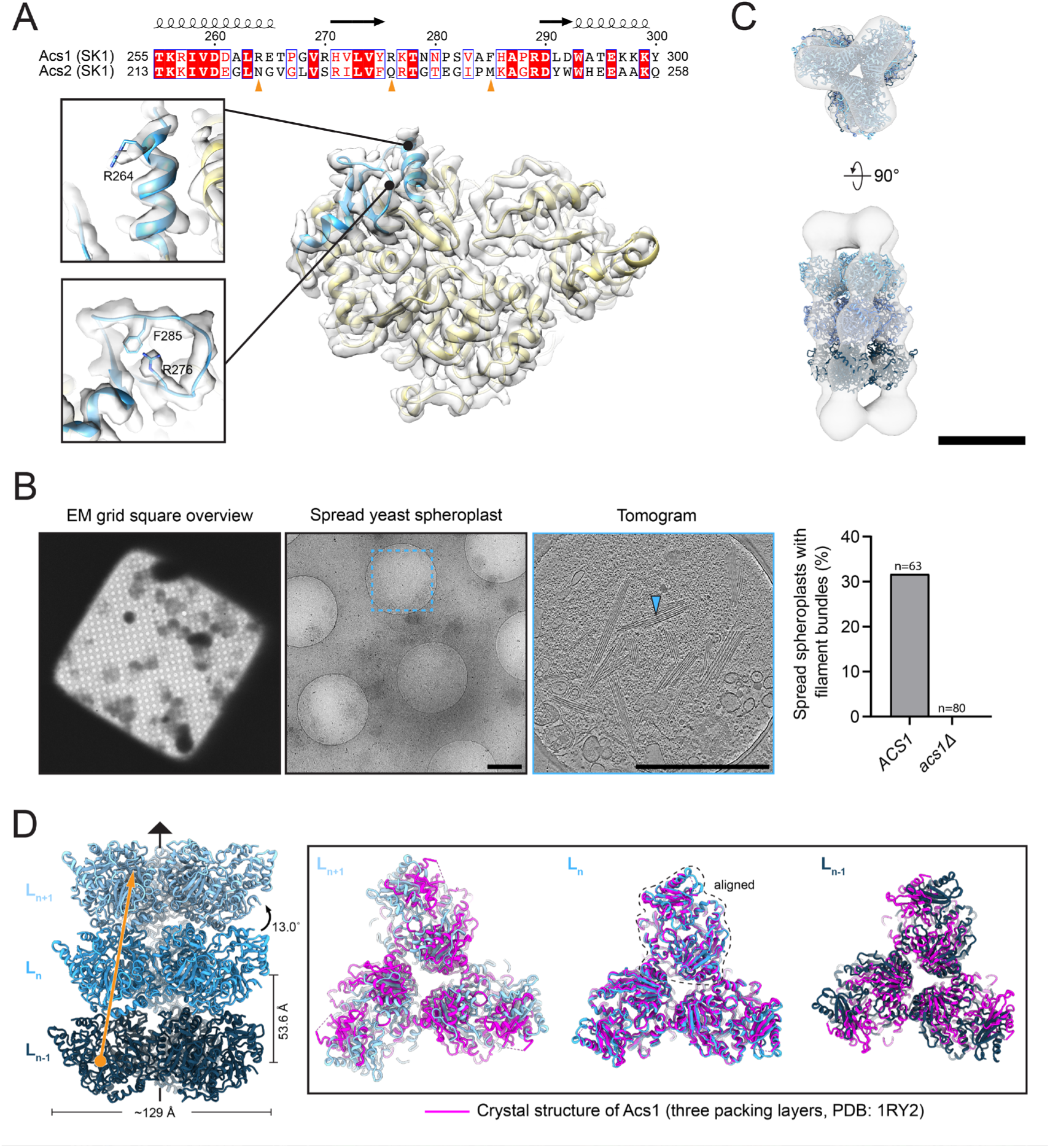
Validation of Acs1 protein identity. **(A)** Reconstructed map unambiguously identifies the Acs1 trimer as the only filament component. Top: sequence alignments of Acs1 and its isoform Acs2 from the SK1 strain. The identical residues, similar residues, and the conserved positions are shown in the same style as in Figure S4D. The secondary structure of Acs1 is shown above the corresponding sequences. Three distinguishing residues of Acs1 (R264, R276, and F285) are highlighted by orange arrowheads and are shown on the bottom panel. Bottom: ribbon and shadowed surface diagrams showing the fitting of Acs1 model with the reconstructed map. The density map is colored gray and is shown in transparent, while the overall model is colored yellow. The aligned parts shown on the top panel are colored blue, while zoom-ins show the densities of three distinguishing residues (R264, R276, and F285). **(B)** Filament bundles consist of Acs1. Left to right: to check for the presence of filament bundles, spread spheroplasts are picked in the EM grid square overview. Afterwards, cryoEM micrographs or tomograms, if needed, are collected to assess whether filament bundles are present (highlighted with a blue arrowhead). Scale bars: 1 µm. Shown are the same images as in Figure S5A. Right: shown are percentages of meiotic spread spheroplasts (6 h after induction of meiosis) containing filament bundles and the total number of spreads (n) analyzed for each genotype are indicated. **(C)** Docking of the high-resolution structure of Acs1 filament fits the sub-tomogram averaged volume of Acs1 filaments from spread yeast spheroplasts as shown in Figure 4B. Three layers of Acs1 trimers are shown in ribbon and colored in different shades of blue as in Figure 4D, while the averaged volume is transparent and colored gray. Scale bar: 10 nm. **(D)** Structural superimposition of Acs1 in the filament assembly and the crystal packing (PDD entry: 1RY2) ^36^. Left: Ribbon diagrams showing three layers of Acs1 trimers, which are colored in different shades of blue as in Figure 4D. The helical parameters (twist = 13.0°, rise = 53.6 Å) are labeled and a single strand is marked with an orange arrowhead. The 3-fold axis is represented by a triangle. Right: Magnified top view of three individual consecutive layers in the Acs1 filament aligned with the crystal packing of yeast Acs1 based on the structural superimposition of the central layer (L_n_). Note that the central layer is nicely aligned, while the neighboring layers (L_n+1_, L_n-1_) mismatch due to the helical twist in the Acs1 filament.

**Figure S8.**
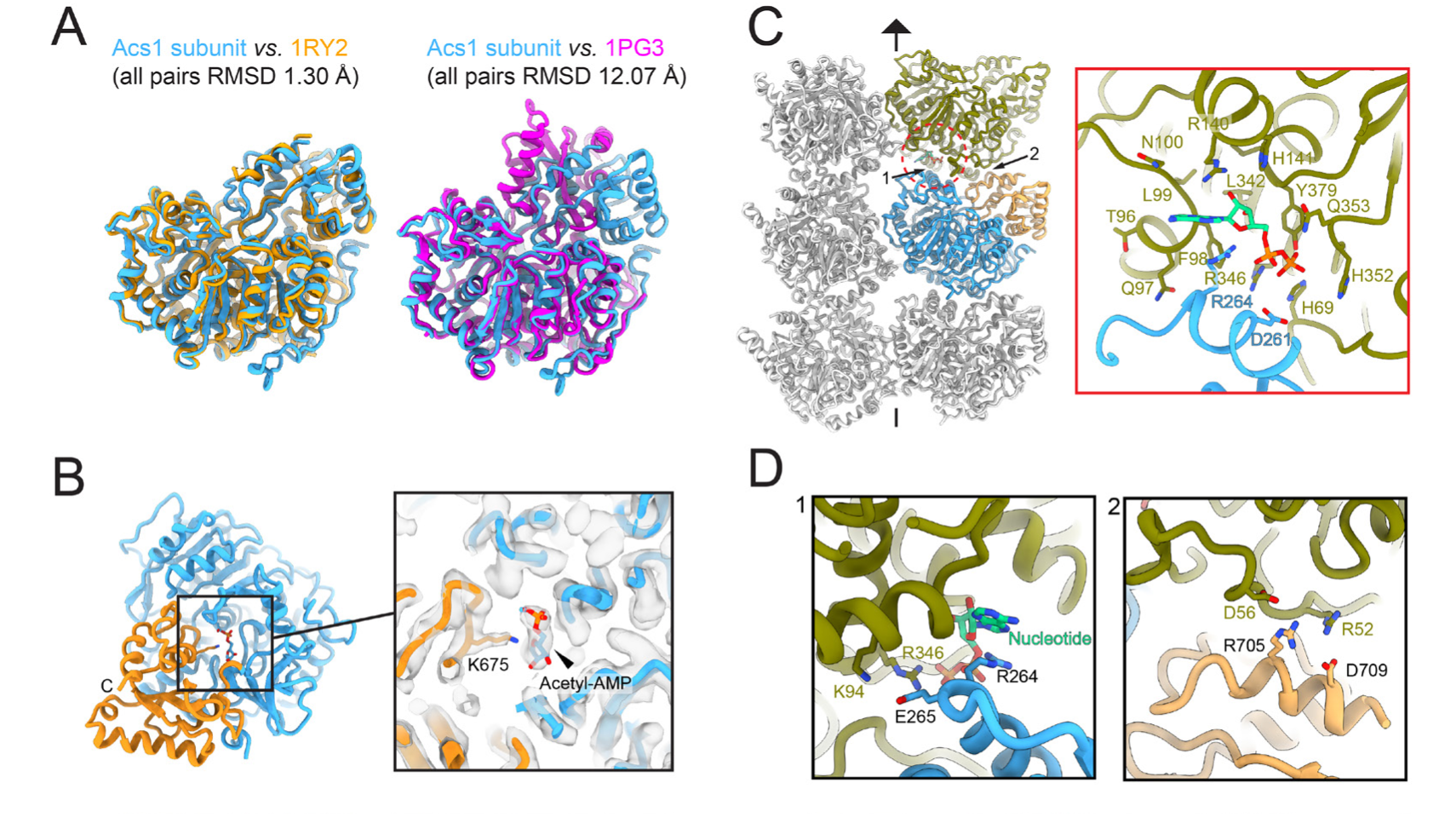
Acs1 subunits bind to two metabolites to form the filament. **(A)** Ribbon diagrams showing that the Acs1 subunit in the filament conformation fits the state that accomplishes the first step of the enzymatic reaction. Structural superpositions show the structural difference between Acs1 subunit from the filament conformation (blue) with the yeast Acs1 binary complex (PDB entry: 1RY2 ^36^, orange, left) representing the first step of the enzymatic reaction, compared to the bacterial ACS homolog ternary complex (PDB entry: 1PG3 ^40^, magenta, right) representing the second step of the enzymatic reaction. **(B)** Ribbon and stick diagrams showing that the acetyl-AMP intermediate interacts with the catalytic residue K675 of Acs1. The Acs1 structure is shown ribbon style and is color-coded as in Figure 5C, whereas the acetyl-AMP intermediate is shown in stick style. The C-terminus of Acs1 is labeled. Zoom-in on the catalytic site is shown on the right, highlighting the contact between the catalytic key residue K675 and the acetyl-AMP intermediate. The density of Acs1 subunit is colored gray and is shown in transparent, while the side chain of K675 is shown in stick style. **(C)** Ribbon and stick diagrams showing the metabolic nucleotide (potentially ADP) bound to two consecutive Acs1 trimer layers. Three Acs1 trimer layers are color-coded as in Figure 5C and the 3-fold axis is represented by a triangle. The binding site of the metabolic nucleotide is highlighted with a red dashed circle and the zoom-in is shown to the right. The metabolic nucleotide and the side chains of the contacting residues are labeled and are shown in stick style. Two interfaces between adjacent Acs1 subunits are marked (1 and 2) and the details are shown in panel (**D**). **(D)** Ribbon and stick diagrams showing that salt bridge pairs contribute to the Acs1 filament assembly. Zoom-ins show the corresponding interfaces highlighted in panel (**C**). The metabolic nucleotide (potentially ADP) and the side chains of residues participating in salt bridge pairs are labeled and are shown in stick style.

**Figure S9.**
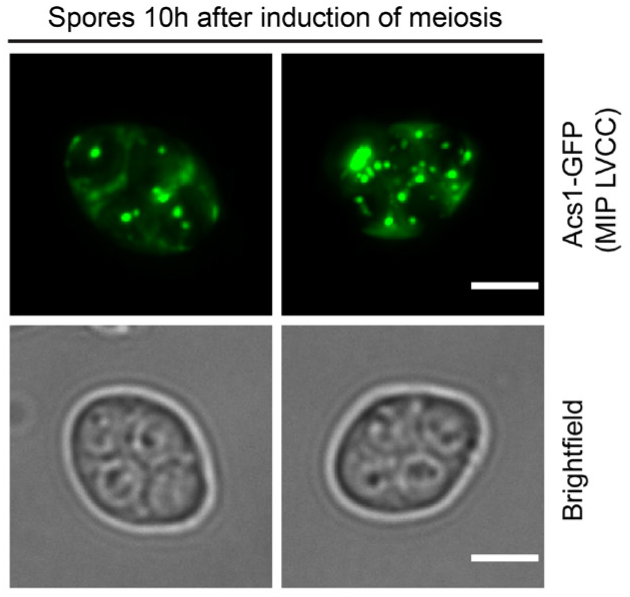
Acs1 C-terminally tagged with GFP forms foci in ascospores. Cells expressing Acs1-GFP from the endogenous promotor were sporulated and imaged by fluorescent light microscopy. Shown are two representative images of maximum intensity z-projections (MIP) after large volume computational clearing (LVCC) and the corresponding Brightfield image of the ascospores. Scale bar: 3 µm.

**Figure S10.**
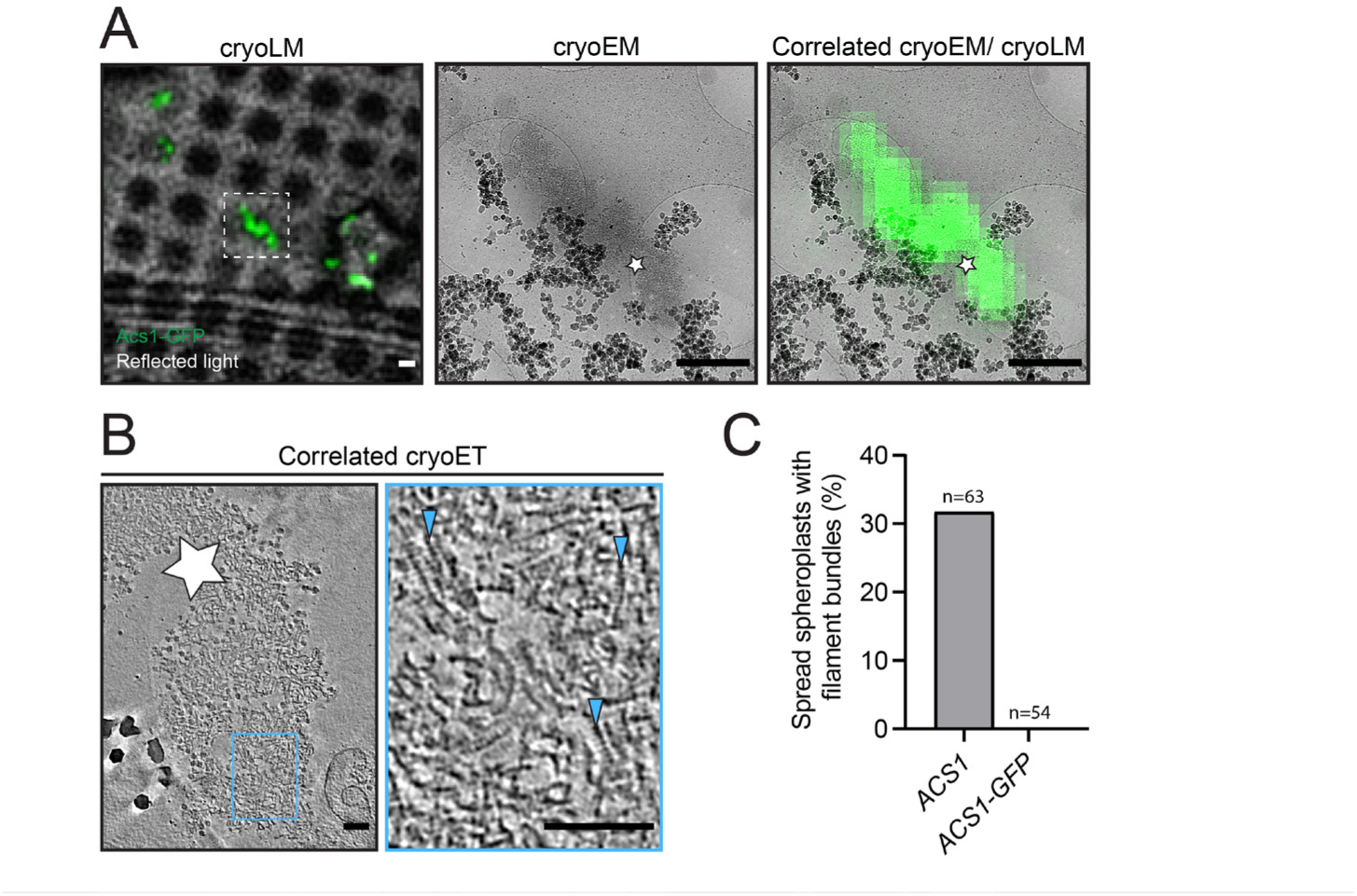
Acs1 C-terminally tagged with GFP forms short single filament aggregates but not elaborate bundles. **(A)** Correlation of Acs1-GFP signal in cryo-confocal light microscopy (cryoLM) with cryoEM. Left: cryoLM of a Acs1-GFP rod from a meiotic spread yeast spheroplast (6 h after induction of meiosis) on an EM grid. Shown is an overlay of maximum intensity projections of Acs1-GFP and reflected light of the EM grid. Middle: the corresponding low resolution EM overview of the area highlighted in the left image. Right: Overlay of the EM overview image with the corresponding Acs1-GFP signal from cryoLM. Scale bars: 1 µm. White star indicates were cryoET imaging was performed for panel (**B**). **(B)** Acs1-GFP forms short ladder-like filament aggregates. Slices through the cryo-tomogram of the corresponding area indicated by white stars in (**A**). Note that short ladder-like filaments, visible in the magnified view (blue box), form an aggregate, which colocalizes with the Acs1-GFP fluorescent signal in (**A**). Single filaments are highlighted with blue arrowheads in the magnified view. Shown are projections of 9.15 nm thick slices. Scale bars: 100 nm. **(C)** No filament bundles are visible in Acs1-GFP spreads. Spread meiotic yeast spheroplasts (6 h after induction of meiosis) for the indicated genotypes were analyzed by cryoEM as described in more detail in Figure S7B. Shown are percentages of spreads containing Acs1 filament bundles in EM images and the total number of spreads (n) analyzed for each genotype are indicated. Note that the experiment was performed together with the strains in Figure S7B. Therefore, the wilde-type *ACS1* strain was re-used in this figure.

**Figure S11.**
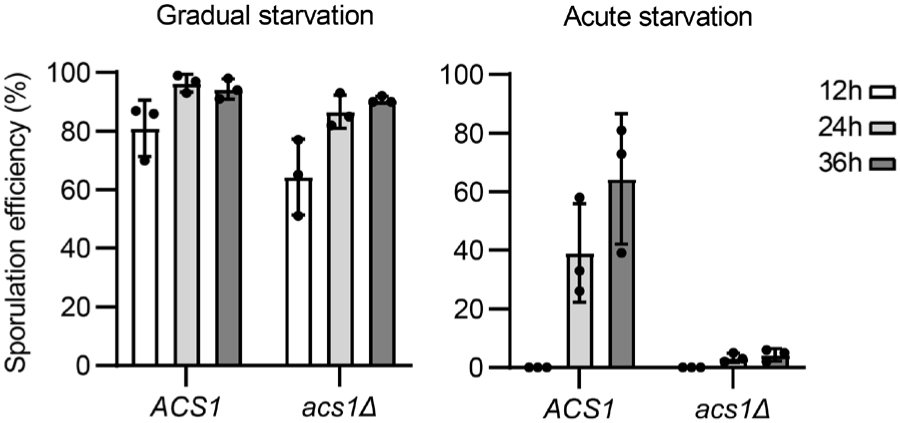
Acs1 is needed for efficient sporulation after acute starvation. Acs1 is needed for efficient sporulation after acute starvation. Yeast strains with the indicated genotypes were induced to enter meiosis by transfer from YPD to SPM (acute starvation), or by transfer from YPD to YPA to SPM (gradual starvation). The efficiency of ascospore formation was quantified by assessing the morphology of 100 cells from three independent experiments after 12, 24 and 36 h in SPM. Ascospore formation was considered successful if at least two spores were enclosed by a spore wall. Plotted values show the mean ± SD.

## SUPPLEMENTARY TABLES

**Table S1:** cryoEM data statistical analysis

|  | Acs1 filament | Ald4 tetramer |
| --- | --- | --- |
| **Data collection and processing** |  |  |
| Nominal magnification | 130,000 | 81,000 |
| Voltage (kV) | 300 |  |
| Electron exposure (e<sup>-</sup>/Å) | ~60 |  |
| Defocus range (μm) | 1.2 - 2.8 |  |
| Pixel size (Å/pixel) | 1.07 (binning = 1) | 1.10 (binning =1) |
| Symmetry imposed | C3 + helical<br>(twist = 13.03°, rise = 53.61 Å) | D2 |
| Final particles (No.) | 17,169 | 29,307 |
| Map resolution (Å) | 3.5 | 3.8 |
| FSC threshold | 0.143 |  |
| **Refinement** |  |  |
| Map sharpening B factor (Å<sup>2</sup>) | -83 | -131 |
| Model composition | (three layers) |  |
| Atoms | 48177 (Hydrogens: 135) | 15416 (Hydrogens: 0) |
| Protein residues | 6084 | 1980 |
| Chains | 18 | 4 |
| Ligands with number | 6R9: 9 | NAP: 4 |
| R.M.S deviations |  |  |
| Bond length (Å) | 0.022 | 0.019 |
| Bond angles (°) | 1.73 | 1.769 |
| Validation |  |  |
| MolProbity score | 1.50 | 1.27 |
| Clashscore | 7.19 | 5.14 |
| Rotamer outlier (%) | 0 | 0 |
| Ramachandran plot |  |  |
| Favored (%) | 97.48 | 98.58 |
| Allowed (%) | 2.52 | 1.42 |
| Outlier (%) | 0 | 0 |
| Masked CC | 0.77 | 0.77 |

**Table S2:**
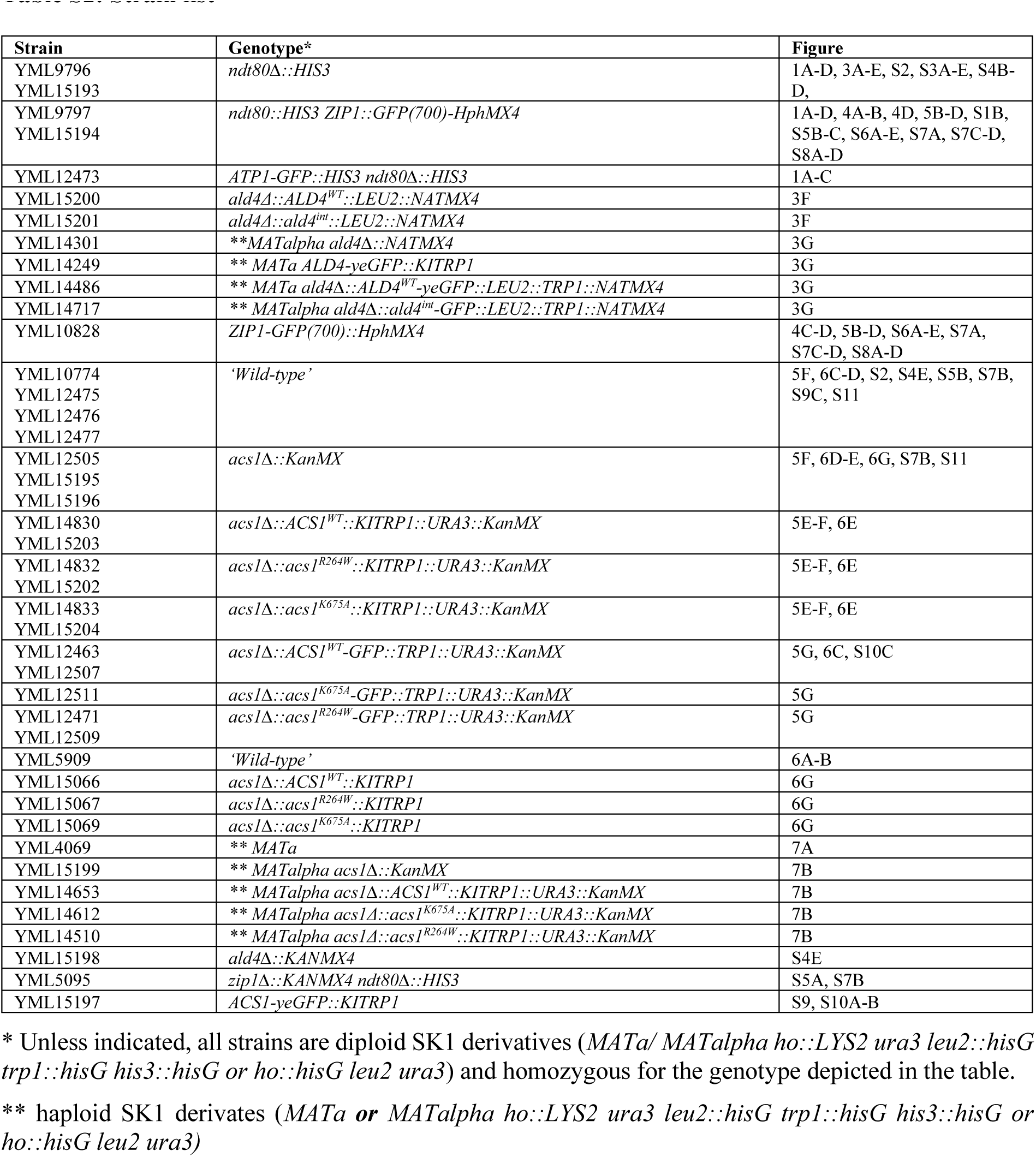
Strain list

**Table S3:** Protein candidates for mitochondrial filaments See separate Excel file

## REFERENCES

1. Lennon, J.T., and Jones, S.E. (2011). Microbial seed banks: the ecological and evolutionary implications of dormancy. Nat Rev Microbiol 9, 119–130. 10.1038/nrmicro2504.

2. Rittershaus, E.S., Baek, S.H., and Sassetti, C.M. (2013). The normalcy of dormancy: common themes in microbial quiescence. Cell Host Microbe 13, 643–651. 10.1016/j.chom.2013.05.012.

3. Aughey, G.N., and Liu, J.L. (2015). Metabolic regulation via enzyme filamentation. Crit Rev Biochem Mol Biol 51, 282–293. 10.3109/10409238.2016.1172555.

4. Garcia-Seisdedos, H., Empereur-Mot, C., Elad, N., and Levy, E.D. (2017). Proteins evolve on the edge of supramolecular self-assembly. Nature 548, 244–247. 10.1038/nature23320.

5. Hvorecny, K.L., and Kollman, J.M. (2023). Greater than the sum of parts: Mechanisms of metabolic regulation by enzyme filaments. Curr Opin Struct Biol 79, 102530. 10.1016/j.sbi.2023.102530.

6. Lynch, E.M., Kollman, J.M., and Webb, B.A. (2020). Filament formation by metabolic enzymes-A new twist on regulation. Curr Opin Cell Biol 66, 28–33. 10.1016/j.ceb.2020.04.006.

7. O’Connell, J.D., Zhao, A., Ellington, A.D., and Marcotte, E.M. (2012). Dynamic reorganization of metabolic enzymes into intracellular bodies. Annu Rev Cell Dev Biol 28, 89–111. 10.1146/annurev-cellbio-101011-155841.

8. Park, C.K., and Horton, N.C. (2019). Structures, functions, and mechanisms of filament forming enzymes: a renaissance of enzyme filamentation. Biophysical Reviews 11, 927–994. 10.1007/s12551-019-00602-6.

9. Simonet, J.C., Burrell, A.L., Kollman, J.M., and Peterson, J.R. (2020). Freedom of assembly: metabolic enzymes come together. Mol Biol Cell 31, 1201–1205. 10.1091/mbc.E18-10-0675.

10. Noree, C., Begovich, K., Samilo, D., Broyer, R., Monfort, E., and Wilhelm, J.E. (2019). A quantitative screen for metabolic enzyme structures reveals patterns of assembly across the yeast metabolic network. Molecular Biology of the Cell 30, 2721–2736. 10.1091/mbc.E19-04-0224.

11. Narayanaswamy, R., Levy, M., Tsechansky, M., Stovall, G.M., O’Connell, J.D., Mirrielees, J., Ellington, A.D., and Marcotte, E.M. (2009). Widespread reorganization of metabolic enzymes into reversible assemblies upon nutrient starvation. Proc Natl Acad Sci U S A 106, 10147–10152. 10.1073/pnas.0812771106.

12. Noree, C., Sato, B.K., Broyer, R.M., and Wilhelm, J.E. (2010). Identification of novel filament-forming proteins in Saccharomyces cerevisiae and Drosophila melanogaster. J Cell Biol 190, 541–551. 10.1083/jcb.201003001.

13. Shen, Q.J., Kassim, H., Huang, Y., Li, H., Zhang, J., Li, G., Wang, P.Y., Yan, J., Ye, F., and Liu, J.L. (2016). Filamentation of Metabolic Enzymes in Saccharomyces cerevisiae. J Genet Genomics 43, 393–404. 10.1016/j.jgg.2016.03.008.

14. Noree, C., Monfort, E., Shiau, A.K., and Wilhelm, J.E. (2014). Common regulatory control of CTP synthase enzyme activity and filament formation. Mol Biol Cell 25, 2282–2290. 10.1091/mbc.E14-04-0912.

15. Hansen, J.M., Horowitz, A., Lynch, E.M., Farrell, D.P., Quispe, J., DiMaio, F., and Kollman, J.M. (2021). Cryo-EM structures of CTP synthase filaments reveal mechanism of pH-sensitive assembly during budding yeast starvation. Elife 10. 10.7554/eLife.73368.

16. Petrovska, I., Nüske, E., Munder, M.C., Kulasegaran, G., Malinovska, L., Kroschwald, S., Richter, D., Fahmy, K., Gibson, K., Verbavatz, J.M., and Alberti, S. (2014). Filament formation by metabolic enzymes is a specific adaptation to an advanced state of cellular starvation. Elife 3. 10.7554/eLife.02409.

17. Nüske, E., Marini, G., Richter, D., Leng, W., Bogdanova, A., Franzmann, T.M., Pigino, G., and Alberti, S. (2020). Filament formation by the translation factor eIF2B regulates protein synthesis in starved cells. Biol Open 9. 10.1242/bio.046391.

18. Stoddard, P.R., Lynch, E.M., Farrell, D.P., Dosey, A.M., DiMaio, F., Williams, T.A., Kollman, J.M., Murray, A.W., and Garner, E.C. (2020). Polymerization in the actin ATPase clan regulates hexokinase activity in yeast. Science 367, 1039–1042. 10.1126/science.aay5359.

19. Neiman, A.M. (2011). Sporulation in the budding yeast Saccharomyces cerevisiae. Genetics 189, 737–765. 10.1534/genetics.111.127126.

20. Marston, A.L., and Amon, A. (2004). Meiosis: cell-cycle controls shuffle and deal. Nat Rev Mol Cell Biol 5, 983–997. 10.1038/nrm1526.

21. Beck, M., and Baumeister, W. (2016). Cryo-Electron Tomography: Can it Reveal the Molecular Sociology of Cells in Atomic Detail? Trends Cell Biol 26, 825–837. 10.1016/j.tcb.2016.08.006.

22. Young, L.N., and Villa, E. (2023). Bringing Structure to Cell Biology with Cryo-Electron Tomography. Annual Review of Biophysics 52, 573–595. 10.1146/annurev-biophys-111622-091327.

23. Oelschlaegel, T., Schwickart, M., Matos, J., Bogdanova, A., Camasses, A., Havlis, J., Shevchenko, A., and Zachariae, W. (2005). The yeast APC/C subunit Mnd2 prevents premature sister chromatid separation triggered by the meiosis-specific APC/C-Ama1. Cell 120, 773–788. 10.1016/j.cell.2005.01.032.

24. Petronczki, M., Matos, J., Mori, S., Gregan, J., Bogdanova, A., Schwickart, M., Mechtler, K., Shirahige, K., Zachariae, W., and Nasmyth, K. (2006). Monopolar attachment of sister kinetochores at meiosis I requires casein kinase 1. Cell 126, 1049–1064. 10.1016/j.cell.2006.07.029.

25. Zachs, T., Schertel, A., Medeiros, J., Weiss, G.L., Hugener, J., Matos, J., and Pilhofer, M. (2020). Fully automated, sequential focused ion beam milling for cryo-electron tomography. Elife 9, e52286. 10.7554/eLife.52286.

26. Xu, L., Ajimura, M., Padmore, R., Klein, C., and Kleckner, N. (1995). NDT80, a meiosis-specific gene required for exit from pachytene in Saccharomyces cerevisiae. Mol Cell Biol 15, 6572–6581. 10.1128/mcb.15.12.6572.

27. Loidl, J., Klein, F., and Engebrecht, J. (1997). Chapter 12 Genetic and Morphological Approaches for the Analysis of Meiotic Chromosomes in Yeast. In Methods in Cell Biology, M. Berrios, ed. (Academic Press), pp. 257–285. 10.1016/S0091-679X(08)60882-1.

28. Loidl, J., Nairz, K., and Klein, F. (1991). Meiotic chromosome synapsis in a haploid yeast. Chromosoma 100, 221–228. 10.1007/bf00344155.

29. Jumper, J., Evans, R., Pritzel, A., Green, T., Figurnov, M., Ronneberger, O., Tunyasuvunakool, K., Bates, R., Žídek, A., Potapenko, A., et al. (2021). Highly accurate protein structure prediction with AlphaFold. Nature 596, 583–589. 10.1038/s41586-021-03819-2.

30. Varadi, M., Anyango, S., Deshpande, M., Nair, S., Natassia, C., Yordanova, G., Yuan, D., Stroe, O., Wood, G., Laydon, A., et al. (2022). AlphaFold Protein Structure Database: massively expanding the structural coverage of protein-sequence space with high-accuracy models. Nucleic Acids Research 50, D439–D444. 10.1093/nar/gkab1061.

31. Wettstein, R., Hugener, J., Gillet, L., Hernández-Armenta, Y., Henggeler, A., Xu, J., Van Gerwen, J., Wollweber, F., Arter, M., Aebersold, R., Beltrao, P., Pilhofer, M., and Matos, J. (unpublished). Waves of regulated protein expression and phosphorylation functionally rewire the proteome to drive gametogenesis.

32. Morgenstern, M., Stiller, S.B., Lübbert, P., Peikert, C.D., Dannenmaier, S., Drepper, F., Weill, U., Höß, P., Feuerstein, R., Gebert, M., et al. (2017). Definition of a High-Confidence Mitochondrial Proteome at Quantitative Scale. Cell Rep 19, 2836–2852. 10.1016/j.celrep.2017.06.014.

33. Perez-Miller, S.J., and Hurley, T.D. (2003). Coenzyme Isomerization Is Integral to Catalysis in Aldehyde Dehydrogenase. Biochemistry 42, 7100–7109. 10.1021/bi034182w.

34. Misonou, Y., Kikuchi, M., Sato, H., Inai, T., Kuroiwa, T., Tanaka, K., and Miyakawa, I. (2014). Aldehyde dehydrogenase, Ald4p, is a major component of mitochondrial fluorescent inclusion bodies in the yeast Saccharomyces cerevisiae. Biol Open 3, 387–396. 10.1242/bio.20147138.

35. Holm, L. (2022). Dali server: structural unification of protein families. Nucleic Acids Research 50, W210–W215. 10.1093/nar/gkac387.

36. Jogl, G., and Tong, L. (2004). Crystal Structure of Yeast Acetyl-Coenzyme A Synthetase in Complex with AMP. Biochemistry 43, 1425–1431. 10.1021/bi035911a.

37. Ma, O.X., Chong, W.G., Lee, J.K.E., Cai, S., Siebert, C.A., Howe, A., Zhang, P., Shi, J., Surana, U., and Gan, L. (2022). Cryo-ET detects bundled triple helices but not ladders in meiotic budding yeast. PLoS One 17, e0266035. 10.1371/journal.pone.0266035.

38. Takagi, T., Osumi, M., and Shinohara, A. (2021). Ultrastructural analysis in yeast reveals a meiosis-specific actin-containing nuclear bundle. Commun Biol 4, 1009. 10.1038/s42003-021-02545-9.

39. Starai, V.J., and Escalante-Semerena, J.C. (2004). Acetyl-coenzyme A synthetase (AMP forming). Cellular and Molecular Life Sciences CMLS 61, 2020–2030. 10.1007/s00018-004-3448-x.

40. Gulick, A.M., Starai, V.J., Horswill, A.R., Homick, K.M., and Escalante-Semerena, J.C. (2003). The 1.75 Å Crystal Structure of Acetyl-CoA Synthetase Bound to Adenosine-5‘-propylphosphate and Coenzyme A. Biochemistry 42, 2866–2873. 10.1021/bi0271603.

41. Starai, V.J., Celic, I., Cole, R.N., Boeke, J.D., and Escalante-Semerena, J.C. (2002). Sir2-dependent activation of acetyl-CoA synthetase by deacetylation of active lysine. Science 298, 2390–2392. 10.1126/science.1077650.

42. Van den Berg, M.A., and Steensma, H.Y. (1995). ACS2, a Saccharomyces cerevisiae gene encoding acetyl-coenzyme A synthetase, essential for growth on glucose. Eur J Biochem 231, 704–713. 10.1111/j.1432-1033.1995.tb20751.x.

43. Takahashi, H., McCaffery, J.M., Irizarry, R.A., and Boeke, J.D. (2006). Nucleocytosolic Acetyl-Coenzyme A Synthetase Is Required for Histone Acetylation and Global Transcription. Molecular Cell 23, 207–217. 10.1016/j.molcel.2006.05.040.

44. Mews, P., Donahue, G., Drake, A.M., Luczak, V., Abel, T., and Berger, S.L. (2017). Acetyl-CoA synthetase regulates histone acetylation and hippocampal memory. Nature 546, 381–386. 10.1038/nature22405.

45. Vasiliou, V., Pappa, A., and Petersen, D.R. (2000). Role of aldehyde dehydrogenases in endogenous and xenobiotic metabolism. Chemico-Biological Interactions 129, 1–19. 10.1016/S0009-2797(00)00211-8.

46. Schug, Z.T., Peck, B., Jones, D.T., Zhang, Q., Grosskurth, S., Alam, I.S., Goodwin, L.M., Smethurst, E., Mason, S., Blyth, K., et al. (2015). Acetyl-CoA synthetase 2 promotes acetate utilization and maintains cancer cell growth under metabolic stress. Cancer Cell 27, 57–71. 10.1016/j.ccell.2014.12.002.

47. Shortall, K., Djeghader, A., Magner, E., and Soulimane, T. (2021). Insights into Aldehyde Dehydrogenase Enzymes: A Structural Perspective. Frontiers in Molecular Biosciences 8. 10.3389/fmolb.2021.659550.

48. Kratzer, S., and Schuller, H.J. (1997). Transcriptional control of the yeast acetyl-CoA synthetase gene, ACS1, by the positive regulators CAT8 and ADR1 and the pleiotropic repressor UME6. Molecular microbiology 26, 631–641. 10.1046/j.1365-2958.1997.5611937.x.

49. Jacobson, M.K., and Bernofsky, C. (1974). Mitochondrial acetaldehyde dehydrogenase from Saccharomyces cerevisiae. Biochimica et biophysica acta 350, 277–291. 10.1016/0005-2744(74)90502-6.

50. Munder, M.C., Midtvedt, D., Franzmann, T., Nüske, E., Otto, O., Herbig, M., Ulbricht, E., Müller, P., Taubenberger, A., Maharana, S., et al. (2016). A pH-driven transition of the cytoplasm from a fluid-to a solid-like state promotes entry into dormancy. Elife 5, e09347. 10.7554/eLife.09347.

51. Joyner, R.P., Tang, J.H., Helenius, J., Dultz, E., Brune, C., Holt, L.J., Huet, S., Müller, D.J., and Weis, K. (2016). A glucose-starvation response regulates the diffusion of macromolecules. Elife 5, e09376. 10.7554/eLife.09376.

52. Walther, T., Létisse, F., Peyriga, L., Alkim, C., Liu, Y., Lardenois, A., Martin-Yken, H., Portais, J.C., Primig, M., and François, J. (2014). Developmental stage dependent metabolic regulation during meiotic differentiation in budding yeast. BMC Biol 12, 60. 10.1186/s12915-014-0060-x.

53. Ray, D., and Ye, P. (2013). Characterization of the metabolic requirements in yeast meiosis. PloS one 8, e63707. 10.1371/journal.pone.0063707.

54. Shvedunova, M., and Akhtar, A. (2022). Modulation of cellular processes by histone and non-histone protein acetylation. Nature Reviews Molecular Cell Biology 23, 329–349. 10.1038/s41580-021-00441-y.

55. Wang, F., Zhang, R., Feng, W., Tsuchiya, D., Ballew, O., Li, J., Denic, V., and Lacefield, S. (2020). Autophagy of an Amyloid-like Translational Repressor Regulates Meiotic Exit. Dev Cell 52, 141–151 e145. 10.1016/j.devcel.2019.12.017.

56. O’Reilly, F.J., Xue, L., Graziadei, A., Sinn, L., Lenz, S., Tegunov, D., Blötz, C., Singh, N., Hagen, W.J.H., Cramer, P., et al. (2020). In-cell architecture of an actively transcribing-translating expressome. Science 369, 554–557. 10.1126/science.abb3758.

57. Xue, L., Lenz, S., Zimmermann-Kogadeeva, M., Tegunov, D., Cramer, P., Bork, P., Rappsilber, J., and Mahamid, J. (2022). Visualizing translation dynamics at atomic detail inside a bacterial cell. Nature 610, 205–211. 10.1038/s41586-022-05255-2.

58. Hoffmann, P.C., Kreysing, J.P., Khusainov, I., Tuijtel, M.W., Welsch, S., and Beck, M. (2022). Structures of the eukaryotic ribosome and its translational states in situ. Nature Communications 13, 7435. 10.1038/s41467-022-34997-w.

59. Xing, H., Taniguchi, R., Khusainov, I., Kreysing, J.P., Welsch, S., Turoňová, B., and Beck, M. (2023). Translation dynamics in human cells visualized at high resolution reveal cancer drug action. Science 381, 70–75. 10.1126/science.adh1411.

60. Ho, C.M., Li, X., Lai, M., Terwilliger, T.C., Beck, J.R., Wohlschlegel, J., Goldberg, D.E., Fitzpatrick, A.W.P., and Zhou, Z.H. (2020). Bottom-up structural proteomics: cryoEM of protein complexes enriched from the cellular milieu. Nat Methods 17, 79–85. 10.1038/s41592-019-0637-y.

61. Skalidis, I., Kyrilis, F.L., Tüting, C., Hamdi, F., Chojnowski, G., and Kastritis, P.L. (2022). Cryo-EM and artificial intelligence visualize endogenous protein community members. Structure 30, 575–589.e576. 10.1016/j.str.2022.01.001.

62. Tüting, C., Kyrilis, F.L., Müller, J., Sorokina, M., Skalidis, I., Hamdi, F., Sadian, Y., and Kastritis, P.L. (2021). Cryo-EM snapshots of a native lysate provide structural insights into a metabolon-embedded transacetylase reaction. Nat Commun 12, 6933. 10.1038/s41467-021-27287-4.

63. Matos, J., Lipp, J.J., Bogdanova, A., Guillot, S., Okaz, E., Junqueira, M., Shevchenko, A., and Zachariae, W. (2008). Dbf4-dependent CDC7 kinase links DNA replication to the segregation of homologous chromosomes in meiosis I. Cell 135, 662–678. S0092-8674(08)01363-9 [pii] 10.1016/j.cell.2008.10.026.

64. Scherthan, H., Wang, H., Adelfalk, C., White, E.J., Cowan, C., Cande, W.Z., and Kaback, D.B. (2007). Chromosome mobility during meiotic prophase in Saccharomyces cerevisiae. Proc Natl Acad Sci U S A 104, 16934–16939. 10.1073/pnas.0704860104.

65. Giaever, G., Chu, A.M., Ni, L., Connelly, C., Riles, L., Véronneau, S., Dow, S., Lucau-Danila, A., Anderson, K., André, B., et al. (2002). Functional profiling of the Saccharomyces cerevisiae genome. Nature 418, 387–391. 10.1038/nature00935.

66. Knop, M., Siegers, K., Pereira, G., Zachariae, W., Winsor, B., Nasmyth, K., and Schiebel, E. (1999). Epitope tagging of yeast genes using a PCR-based strategy: more tags and improved practical routines. Yeast 15, 963–972. 10.1002/(sici)1097-0061(199907)15:10b<963::Aid-yea399>3.0.Co;2-w.

67. Schindelin, J., Arganda-Carreras, I., Frise, E., Kaynig, V., Longair, M., Pietzsch, T., Preibisch, S., Rueden, C., Saalfeld, S., Schmid, B., et al. (2012). Fiji: an open-source platform for biological-image analysis. Nat Methods 9, 676–682. 10.1038/nmeth.2019.

68. Wild, P., Susperregui, A., Piazza, I., Dörig, C., Oke, A., Arter, M., Yamaguchi, M., Hilditch, A.T., Vuina, K., Chan, K.C., et al. (2019). Network Rewiring of Homologous Recombination Enzymes during Mitotic Proliferation and Meiosis. Mol Cell 75, 859–874.e854. 10.1016/j.molcel.2019.06.022.

69. Salah, S.-M., and Nasmyth, K. (2000). Destruction of the securin Pds1p occurs at the onset of anaphase during both meiotic divisions in yeast. Chromosoma 109, 27–34. 10.1007/s004120050409.

70. Matos, J., Blanco, Miguel G., Maslen, S., Skehel, J.M., and West, Stephen C. (2011). Regulatory Control of the Resolution of DNA Recombination Intermediates during Meiosis and Mitosis. Cell 147, 158–172. 10.1016/j.cell.2011.08.032.

71. Faelber, K., Dietrich, L., Noel, J.K., Wollweber, F., Pfitzner, A.K., Mühleip, A., Sánchez, R., Kudryashev, M., Chiaruttini, N., Lilie, H., et al. (2019). Structure and assembly of the mitochondrial membrane remodelling GTPase Mgm1. Nature 571, 429–433. 10.1038/s41586-019-1372-3.

72. Iancu, C.V., Tivol, W.F., Schooler, J.B., Dias, D.P., Henderson, G.P., Murphy, G.E., Wright, E.R., Li, Z., Yu, Z., Briegel, A., et al. (2006). Electron cryotomography sample preparation using the Vitrobot. Nat Protoc 1, 2813–2819. 10.1038/nprot.2006.432.

73. Tivol, W.F., Briegel, A., and Jensen, G.J. (2008). An improved cryogen for plunge freezing. Microsc Microanal 14, 375–379. 10.1017/S1431927608080781.

74. Schorb, M., and Briggs, J.A.G. (2014). Correlated cryo-fluorescence and cryo-electron microscopy with high spatial precision and improved sensitivity. Ultramicroscopy 143, 24–32. 10.1016/j.ultramic.2013.10.015.

75. Mastronarde, D.N. (2005). Automated electron microscope tomography using robust prediction of specimen movements. J Struct Biol 152, 36–51. 10.1016/j.jsb.2005.07.007.

76. Kremer, J.R., Mastronarde, D.N., and McIntosh, J.R. (1996). Computer visualization of three-dimensional image data using IMOD. J Struct Biol 116, 71–76. 10.1006/jsbi.1996.0013.

77. Tegunov, D., and Cramer, P. (2019). Real-time cryo-electron microscopy data preprocessing with Warp. Nature Methods 16, 1146–1152. 10.1038/s41592-019-0580-y.

78. Heebner, J.E., Purnell, C., Hylton, R.K., Marsh, M., Grillo, M.A., and Swulius, M.T. (2022). Deep Learning-Based Segmentation of Cryo-Electron Tomograms. J Vis Exp. 10.3791/64435.

79. Pettersen, E.F., Goddard, T.D., Huang, C.C., Meng, E.C., Couch, G.S., Croll, T.I., Morris, J.H., and Ferrin, T.E. (2021). UCSF ChimeraX: Structure visualization for researchers, educators, and developers. Protein Sci 30, 70–82. 10.1002/pro.3943.

80. Castaño-Díez, D., Kudryashev, M., Arheit, M., and Stahlberg, H. (2012). Dynamo: a flexible, user-friendly development tool for subtomogram averaging of cryo-EM data in high-performance computing environments. J Struct Biol 178, 139–151. 10.1016/j.jsb.2011.12.017.

81. Rosenthal, P.B., and Henderson, R. (2003). Optimal determination of particle orientation, absolute hand, and contrast loss in single-particle electron cryomicroscopy. J Mol Biol 333, 721–745. 10.1016/j.jmb.2003.07.013.

82. Scheres, S.H.W. (2012). RELION: Implementation of a Bayesian approach to cryo-EM structure determination. Journal of Structural Biology 180, 519–530. 10.1016/j.jsb.2012.09.006.

83. Zheng, S.Q., Palovcak, E., Armache, J.-P., Verba, K.A., Cheng, Y., and Agard, D.A. (2017). MotionCor2: anisotropic correction of beam-induced motion for improved cryo-electron microscopy. Nature Methods 14, 331–332. 10.1038/nmeth.4193.

84. Zhang, K. (2016). Gctf: Real-time CTF determination and correction. J Struct Biol 193, 1–12. 10.1016/j.jsb.2015.11.003.

85. Zivanov, J., Nakane, T., Forsberg, B.O., Kimanius, D., Hagen, W.J.H., Lindahl, E., and Scheres, S.H.W. (2018). New tools for automated high-resolution cryo-EM structure determination in RELION-3. Elife 7, e42166. 10.7554/eLife.42166.

86. He, S., and Scheres, S.H.W. (2017). Helical reconstruction in RELION. Journal of Structural Biology 198, 163–176. 10.1016/j.jsb.2017.02.003.

87. Sanchez-Garcia, R., Gomez-Blanco, J., Cuervo, A., Carazo, J.M., Sorzano, C.O.S., and Vargas, J. (2021). DeepEMhancer: a deep learning solution for cryo-EM volume post-processing. Communications Biology 4, 874. 10.1038/s42003-021-02399-1.

88. Emsley, P., Lohkamp, B., Scott, W.G., and Cowtan, K. (2010). Features and development of Coot. Acta Crystallogr D Biol Crystallogr 66, 486–501. 10.1107/s0907444910007493.

89. Song, Y., DiMaio, F., Wang, R.Y., Kim, D., Miles, C., Brunette, T., Thompson, J., and Baker, D. (2013). High-resolution comparative modeling with RosettaCM. Structure 21, 1735–1742. 10.1016/j.str.2013.08.005.

90. Adams, P.D., Afonine, P.V., Bunkóczi, G., Chen, V.B., Davis, I.W., Echols, N., Headd, J.J., Hung, L.W., Kapral, G.J., Grosse-Kunstleve, R.W., et al. (2010). PHENIX: a comprehensive Python-based system for macromolecular structure solution. Acta Crystallogr D Biol Crystallogr 66, 213–221. 10.1107/s0907444909052925.

91. Krissinel, E., and Henrick, K. (2007). Inference of macromolecular assemblies from crystalline state. J Mol Biol 372, 774–797. 10.1016/j.jmb.2007.05.022.

92. Gouet, P., Courcelle, E., Stuart, D.I., and Métoz, F. (1999). ESPript: analysis of multiple sequence alignments in PostScript. Bioinformatics 15, 305–308. 10.1093/bioinformatics/15.4.305.

